# Fibroblasts from metastatic sites induce broad-spectrum drug desensitization via modulation of mitochondrial priming

**DOI:** 10.1101/197376

**Authors:** Benjamin D. Landry, Thomas Leete, Ryan Richards, Peter Cruz-Gordillo, Gary Ren, Alyssa D. Schwartz, Shelly R. Peyton, Michael J. Lee

## Abstract

Due to tumor heterogeneity, most believe that effective treatments should be tailored to the features of an individual tumor or tumor subclass. It is still unclear what information should be considered for optimal disease stratification, and most prior work focuses on tumor genomics. Here, we focus on the tumor micro-environment. Using a large-scale co-culture assay optimized to measure drug-induced cell death, we identify tumor-stroma interactions that modulate drug sensitivity. Our data show that the chemo-insensitivity typically associated with aggressive subtypes of breast cancer is not cell intrinsic, but rather a product of tumor-fibroblast interactions. Additionally, we find that fibroblast cells influence tumor drug response in two distinct and divergent manners, which were predicable based on the anatomical origin from which the fibroblasts were harvested. These divergent phenotypes result from modulation of “mitochondrial priming” of tumor cells, caused by secretion of inflammatory cytokines, such as IL6 and IL8, from stromal cells.

## INTRODUCTION

DNA damaging agents continue to be used as frontline therapies in the treatment of most forms of cancer. These therapies are effective in many cases; however, sensitivity is extremely variable, even amongst tumor cells of a single stratified subtype (Fry et al., 2008). For instance, “triple-negative” breast cancers (TNBCs) – a subtype defined only by the lack of estrogen and progesterone receptor expression, and lack of HER2 amplification – are the most chemo-sensitive subtype of breast cancer, but also the subtype with the shortest disease-free survival and lowest overall survival rates (Anders and Carey, 2008; Carey et al., 2007). This paradox is thought to result from heterogeneity within the TNBC subclass. Recent studies have highlighted that TNBC is likely not a single disease, but rather an amalgamation of several distinct diseases (Lehmann et al., 2011). Nonetheless, although many tumor subtypes like TNBC are now known to be heterogeneous, it remains unclear which features of this heterogeneity are responsible for creating the variable chemosensitivity that is observed.

The study of tumor heterogeneity has generally focused on the genomics of tumor cells (Cancer Genome Atlas Network, 2012; Cancer Genome Atlas Network et al., 2012; Shah et al., 2012). Several studies now exist that have explored the relationship between tumor genetics or tumor gene expression and drug response (Barretina et al., 2012; Cohen et al., 2011; Lamb et al., 2006; Li et al., 2017). Many insights have been gained from these and other studies, but even collectively, these studies fail to create a clear understanding of the variable levels of sensitivity to commonly used chemotherapeutics (Innocenti et al., 2011; Jiang et al., 2016). An important consideration is that substantial non-genetic heterogeneity exists within tumors, and these influences are generally missed in studies that focus exclusively on tumor genomics. For instance, several classes of normal cells typically reside within tumors, and in some cases these have been demonstrated to alter tumor cell behavior, including sensitivity to drugs (Pallasch et al., 2014).

It is increasingly recognized that many tumor phenotypes, including tumor initiation, epithelial to mesenchymal transition (EMT), and metastatic potential, are the influenced by interactions between cancer cells and the fibroblasts residing within or near tumor cells (Kalluri and Zeisberg, 2006). The role of these interactions in drug sensitivity has been explored using *in vitro* co-culture systems, in which cancer cell specific expression of luciferase (Mcmillin et al., 2010) or GFP (Straussman et al., 2012) was used to specifically quantify tumor cell drug sensitivity in the presence or absence of other stromal cell types. These studies revealed that stromal fibroblasts are a common, sometimes potent, modulator of drug sensitivity, generally resulting in de-sensitization or drug resistance. Additionally, these large-scale studies have revealed significant variability in the nature of the tumor-stroma interactions, in which the drug sensitivity of cancer cells appears to depend on the particular combination of tumor cell, stromal cell, and drug used (Mcmillin et al., 2010).

Here, we develop a mixed co-culture assay optimized to specifically quantify cell death rather than cell proliferation, and use this assay to characterize functional interactions between tumor cells, stromal cells, and anti-cancer chemotherapeutic agents. Our work reveals two previously unappreciated principles by which stromal fibroblasts alter the tumor’s drug response. First, our study finds that the drug insensitivity associated with some aggressive subtypes of breast cancer is not a cell intrinsic property, but rather a product of tumor-fibroblast interactions. Second, we find that fibroblast cells influence tumor drug response in two distinct and divergent manners, which were predicable based on the anatomical origin from which the fibroblasts were harvested. Specifically, we found that fibroblasts harvested from locations that are common sites of breast cancer metastasis – bone, liver, lung, or brain – promote broad-spectrum drug resistance. Conversely, fibroblasts from locations that do not typically harbor metastatic breast cancer – such as the uterus or skin – promote broad-spectrum drug sensitization. Mechanistically, these fibroblast-dependent phenotypes result from modulation of mitochondrial “priming”, which changes the threshold for initiation of apoptosis in cancer cells. Furthermore, we find that the interaction between tumor cells and fibroblasts occurs through inflammatory cytokines, such as IL6 and IL8, which are secreted by fibroblasts and potently augment tumor cell drug sensitivity. Taken together, our study highlights the tumor micro-environment as an important source of drug resistance for aggressive breast cancers, and reveals new strategies for sensitizing these cells to conventional chemotherapies.

## RESULTS

### Cell intrinsic sensitivity to commonly used chemotherapy is similar for basal-like and mesenchymal-like TNBC cells

In the use of molecularly targeted therapies, genetic stratification has led to significant improvements in treatment efficacy (Al-Lazikani et al., 2012). In contrast, the use of DNA-damaging agents is not typically informed by genomic or gene expression features, and instead are selected based on the anatomical origin of the disease. Based on prior genetic studies, several mutations have been identified that alter DNA damage sensitivity, but these are generally restricted to those that alter drug availability or mutations that directly affect DNA repair (Innocenti et al., 2011; Yard et al., 2016). To identify a more complete understanding of the molecular features that contribute to variation in DNA damage sensitivity, we focused on TNBC cells, due to the known variation in drug response that is observed within this subclass. TNBCs are treated exclusively with DNA damaging therapies, and responses are variable, even in the absence of mutations known to alter DNA repair (Lehmann et al., 2011).

To highlight this variability in the drug response, we selected a panel of ten TNBC cells from either the “basal-like” or “mesenchymal-like” expression classes (Heiser et al., 2009; Lehmann et al., 2011; Perou et al., 2000). Basal-like (BL) cells – sometimes referred to as “Basal A”, “Basal-like 1”, and “Basal-like 2” – are defined by expression of basal or myoepithelial genes. These cells are highly proliferative, tend to have elevated expression of DNA damage response genes, and generally respond at higher rates to cytotoxic chemotherapies (Lehmann et al., 2011). Mesenchymal-like (ML) TNBCs – which includes “Mesenchymal”, “Mesenchymal stem-like”, and also “claudin-low” expression classes – are enriched for expression of genes related to EMT, and genes associated with stemness. These cells are more “aggressive” clinically, more de-differentiated, more metastatic, and more chemo-resistant *in vivo* (Lehmann et al., 2011; Prat et al., 2010). Thus, we reasoned that identifying mechanisms which account for the variability in DNA damage sensitivity between the BL and ML classes may aid in patient stratification or help to create new strategies for improving responses to these agents.

We first profiled the response of TNBC cells to doxorubicin (also called Adriamycin), a topoisomerase II inhibitor that is commonly given to TNBC patients. Since doxorubicin is given as frontline therapy to all breast cancer patients, we suspected that if the observed clinical patterns of aggressiveness were due to intrinsic differences in drug sensitivity associated with these expression states, different levels of sensitivity to this drug should be observed *in vitro*. Indeed, the least sensitive cells were HCC-1395, a TNBC of the ML expression state; the most sensitive cells were MDA-MB-468, a TNBC in the chemo-sensitive BL category (Figure 1A). In contrast, however, the rest of the cell lines tested were similarly sensitive to doxorubicin, regardless of their gene expression state. To see if this was unique to doxorubicin, we also profiled responses to other Topo I and II inhibitors in this panel of cells. Overall, these data reveal relatively similar levels of drug sensitivity across these 10 cell lines (Supplemental Figure 1A). For example, for most of the drugs tested, the EC_50_ values for at least 8 of the 10 cells were within technical error range of our assay. Furthermore, for all drugs except camptothecin, we failed to observe any obvious separation between BL and ML cells. To more rigorously determine if the patterns of sensitivity to these drugs could be used to distinguish BL versus ML cells, we performed hierarchical clustering using either the EC_50_ or the maximum effect observed for each drug. This analysis also failed to separate these two expression states based on their observed drug sensitivity profile (Figure 1B and Supplemental Figure 1B). Finally, we also analyzed data publically available through the LINCS consortium, which include drug sensitivities for a larger panel of 27 BL or ML cell lines. These data also show that BL and ML cells have similar levels of sensitivity to topoisomerase inhibitors, specifically, or to all anti-cancer drugs, generally (Figure 1C). Thus, our data suggest that the differences in DNA damage chemo-sensitivity that are commonly observed for BL and ML tumors *in vivo* are not an intrinsic property associated with the BL and ML expression states of these cells.

**Figure 1:**
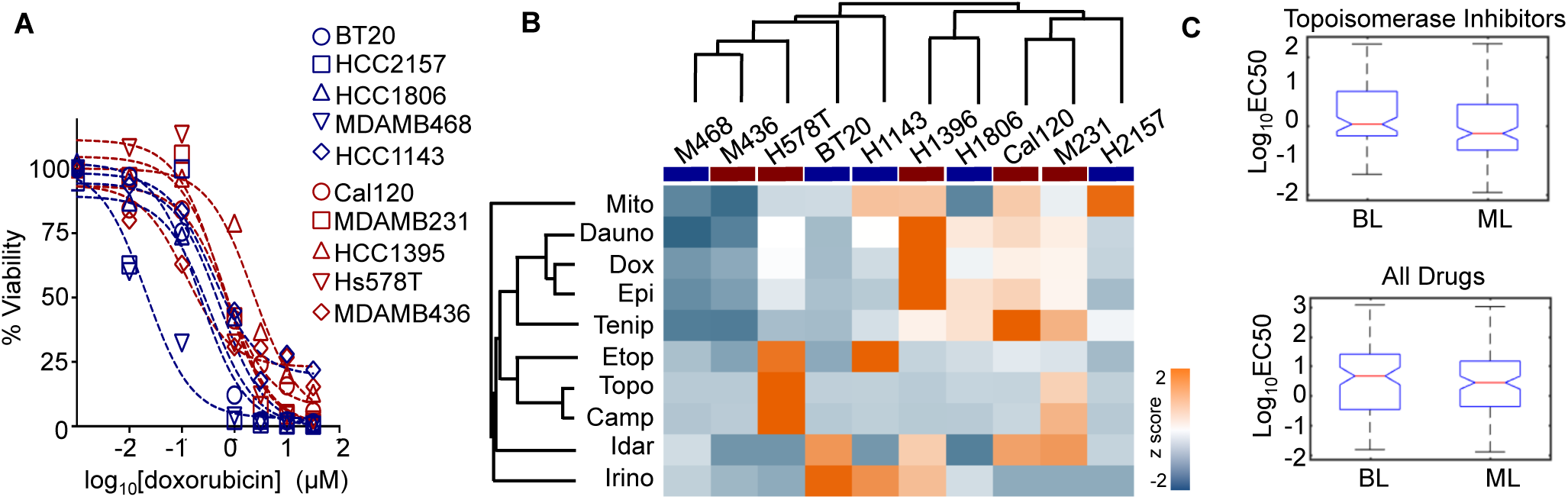
TNBC cell sensitivity to topoisomerase inhibition is not well predicted by basal-like versus mesenchymal-like gene expression status. **(A)** Panel of 10 TNBC cell lines from the Basal-like (BL, blue) or Mesenchymal-like (ML, red) gene expression subclasses. Relative viability following 72 hour exposure to doxorubicin quantified using CellTiter-Glo. Data are from biological duplicates. **(B)** Cell viability measured as in (A) for 10 common Topo I or II inhibitors. Data are z scored EC50 per drug. Dendograms from hierarchical clustering shown for drugs and for cells (BL cells highlighted with blue bar; ML cells highlighted with red bar). **(C)** Sensitivity to topoisomerase inhibitors (top) or all drugs (bottom) in publically available LINCS data. Data are representative of 27 TNBC cell lines and 67 total drugs.

### Co-culture screen to identify environmental influences on drug sensitivity

Based on the results of our *in vitro* drug screen of TNBC cells grown in standard mono-culture conditions, we sought to identify cell non-autonomous sources that may affect sensitivity to DNA damaging agents. Several recent studies have suggested that interactions between tumor cells and components of the tumor micro-environment – including extracellular matrix, growth factors, and other stromal cell types – can alter sensitivity to chemotherapy (Nguyen et al., 2014; Straussman et al., 2012; Weaver et al., 2002). We focused on interactions between cancer cells and stromal fibroblasts, which are often the predominant stromal type found within tumors.Fibroblast infiltration has prognostic and predictive significance in many cancers, generally associated with poor outcomes (Kalluri and Zeisberg, 2006). Moreover, fibroblasts are well known to supply growth factors and matrix proteins, which may alter how cells respond to DNA damage (Lee et al., 2012).

To identify interactions between tumor cells and stromal fibroblasts that alter drug response, we used an *in vitro* co-culture system that was successfully used to study tumor-stroma interactions in other contexts (Straussman et al., 2012). In this experimental platform cancer cells are genetically modified to express GFP, which facilitates rapid, quantitative, high-throughput, and cancer cell specific measurement of drug response dynamics. We piloted this study using BT-20 TNBC cells, either grown in mono-culture or in co-culture with HADF, a primary non-immortalized human fibroblast cell harvested from the adrenal gland. We determined α-smooth muscle actin (SMA) expression, a marker of the “activated fibroblast” expression state, which is commonly observed in fibroblasts associated with tumors (also called Cancer Associated Fibroblasts, or CAFs, and myofibroblasts). Immunofluorescence microscopy experiments confirmed that SMA expression in HADF cells is generally low and variable when these cells are grown in mono-culture. SMA expression is increased in HADF cell co-cultured with BT-20 TNBC cells, and a similar pattern was observed for other primary fibroblasts (Supplemental Figure 2). For our pilot drug screen, these cells were exposed to one of two drugs: erlotinib, a small molecule EGFR inhibitor, or camptothecin, a potent Topo I inhibitor (Supplemental Figure 3A). Erlotinib does not kill BT-20 cells but does induce a transient growth arrest (i.e. cytostasis), whereas camptothecin potently kills BT-20 cells (i.e. cytotoxicity) (Lee et al., 2012). Total well fluorescence measured using a fluorescence plate reader revealed that co-culture with HADF enhanced the proliferation rate of BT-20 cells to a small extent. Furthermore, we found that co-culture with HADF potently blocked erlotinib-mediated cytostasis of BT-20 cells, but had no effect on camptothecin sensitivity. Notably, well-based fluorescence failed to capture the potent death that we observe by other methods following camptothecin exposure, with all measurements in the time course recording higher values than the initial pre-drug measurement (Supplemental Figure 3A and 1A). We were concerned that this reflected a poor sensitivity of this assay, particularly with respect to quantifying the degree of cytotoxicity rather than proliferation. Thus, 96 hours after drug exposure, we collected images of these wells using fluorescence microscopy and quantified cell numbers from these images using a CellProfiler-based automated analysis pipeline (Lamprecht et al., 2007). Our quantitative image analysis confirmed the growth rate increase induced by HADF, as well as the loss of erlotinib-induced cytostasis in HADF co-culture (Supplemental Figure 3B). Importantly, however, image analysis revealed a strong stromal interaction that was not observed by well-based fluorescence measurements. Less than 1% of BT-20 cells survived chronic camptothecin exposure if grown in mono-culture, but roughly 25% of these cells survived in the presence of HADF (Supplemental Figure 3C). Taken together, these data indicate that well-based measurement of GFP fluorescence is appropriate for quantifying changes to proliferation, but not sufficient for quantifying the degree of cell death in a population of cells.

### Co-culture screen optimized to monitor cytotoxicity reveals widespread stromal influence on TNBC drug sensitivity

Because we were primarily focused on the study of cytotoxic DNA damaging agents, we aimed to modify our co-culture screen design to optimize measurement of cell death. We used JC- 1, a dye that accumulates within mitochondria and is often used as a surrogate measure of apoptotic cell death (Figure 2A and B) (Montero et al., 2015). At low concentrations, this dye exists as a monomer and yields green fluorescence, however, when accumulated at high concentrations within mitochondria, this dye forms aggregates, which yield orange/red fluorescence. Thus, the red fluorescence of the JC-1 dye reports cellular mitochondrial integrity, which is lost when cells activate apoptosis. To assess the suitability of JC-1 to quantify modulation in the degree of cell death, we again piloted this assay on BT-20 cells treated with camptothecin in the presence or absence of HADF. Images of these cells taken prior to drug exposure confirm punctate red fluorescence in BT-20, but not HADF, confirming that the dye is not exchanged between cells in co-culture (Figure 2B). 96 hours after exposure to camptothecin, the majority of BT-20 cells had significantly reduced JC-1 red fluorescence, suggesting that mitochondrial integrity has been compromised (Figure 2C). Importantly, JC-1 red fluorescence measured using a fluorescence plate reader was sufficiently sensitive for observing both the potent cell death of BT-20 cells in mono-culture, and the protective effect of HADF cells in co-culture (Figure 2D).

**Figure 2:**
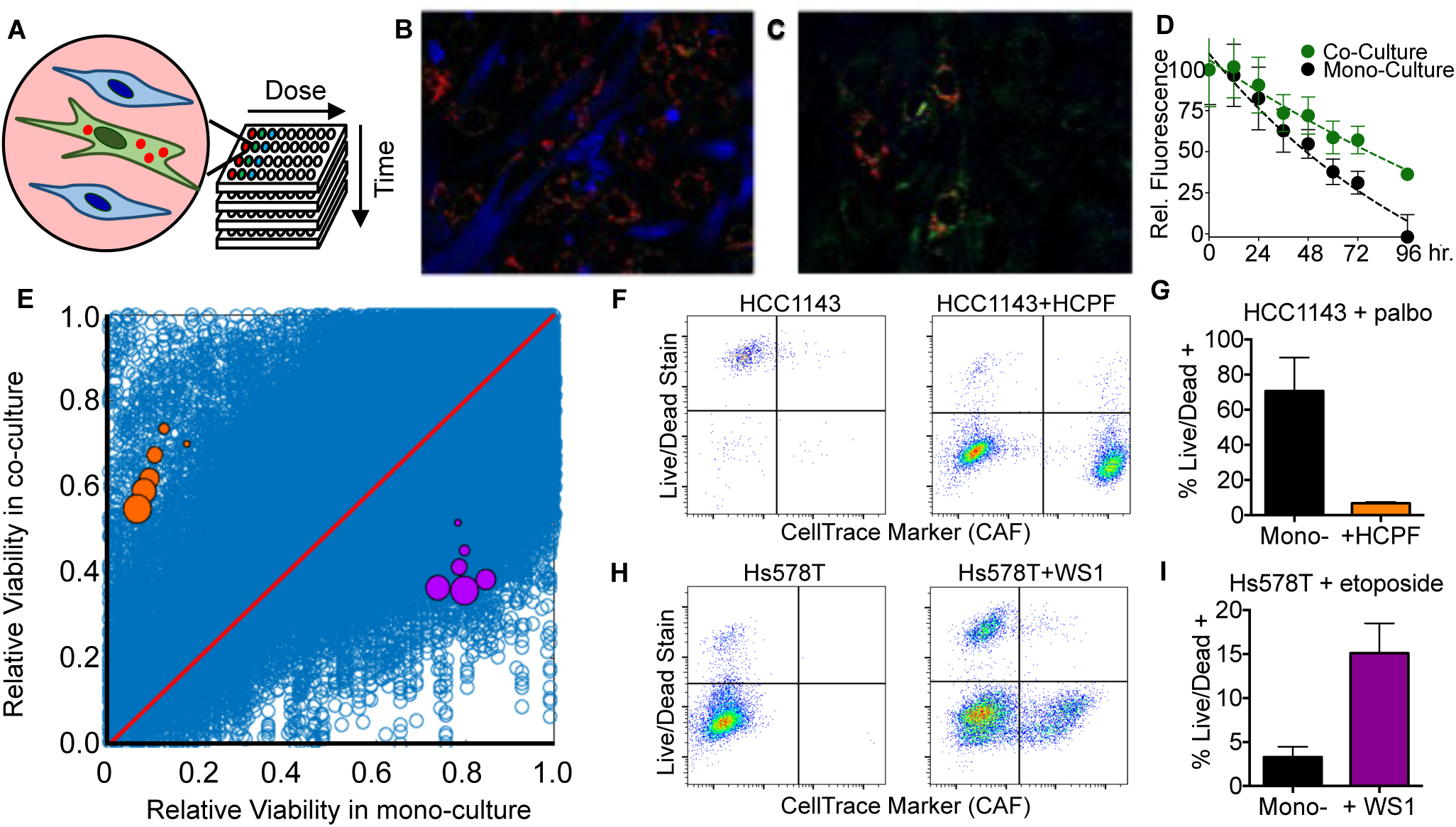
Co-culture screen to identify tumor-stroma interactions that modulate drug induced cell death. **(A)** Schematic of screen design. TNBC cell lines were labeled with JC-1,grown in mono-culture or in co-culture with primary fibroblast cells, and treated with one of 42 anti-cancer drugs. JC-1 fluorescence was monitored using a fluorescence plate reader at 8 hour intervals for 72 hours. **(B and C)** Representative images of BT20 cells co-cultured with HADF fibroblasts. BT20 cells labeled with JC-1 dye; HADF labeled with a blue cell dye (CellTrace). Images taken before drug addition (B) or 96 hours after exposure to 0.5 µM camptothecin (C). (D) Kinetic trace of JC-1 red fluorescence following exposure to camptothecin as in panel B-C. Data are relative JC-1 red fluorescence, compared to well average prior to drug addition. (E) Total co-culture screen data. 312,120 total measurements of drug response. Orange dots represent HCC1143 cells co-cultured with HCPF and treated with palbociclib. Purple dots represent Hs578T co-cultured with WS1 and exposed to etoposide. For colored dots, increasing size represents longer drug exposure times. (F-I) Validation of co-culture screening data. Error bars represent mean +/- standard deviation for five (panel D) and three (G,I) biological replicates.

To evaluate the role of stromal fibroblasts in DNA damage sensitivity, we selected six TNBC cell lines (3 BL and 3 ML) that have relatively similar levels of sensitivity to DNA damage. These JC-1 labeled TNBC cells were grown in mono-culture or in co-culture with each of a panel of 16 primary human fibroblasts. Each culture was exposed to increasing doses of 42 anticancer drugs (at least one drug per class for all current FDA approved breast cancer drugs, Supplemental Tables 1 and 2). JC-1 red fluorescence was quantified at 8 hour intervals for 72 hours. In total, we collected more than 300,000 measurements of drug sensitivity (Figure 2E and Supplemental Table 3). We found a strong overall correlation among biological replicates, indicating that the stromal influences observed were not due to measurement noise (Supplemental Figure 4). To identify TNBC-fibroblast interactions that significantly altered sensitivity, we used a statistical fold-change cut-off of 3x the standard deviation observed among replicates. This analysis identified 5039 significantly changed drug responses (Supplemental Figure 4D). This list of “hits” was significantly depleted for responses at early times (i.e. 8 hours), low doses (0.1 µM), and responses to anti-estrogen drugs (Supplemental Table 4). Non-response to anti-estrogen compounds is expected as TNBCs do not express estrogen or progesterone receptors.

The majority of TNBC cell-fibroblast interactions did not alter drug sensitivity (Supplemental Figure 4A-B). Nonetheless, our screen revealed many striking phenotypes, which strongly altered drug sensitivity in both positive and negative directions. To determine the reliability of these measurements, we selected both strong and moderate phenotypes to validate by flow cytometry. For example, our screen identified that palbociclib killed more than 80% of HCC-1143 cells, a Basal-Like TNBC, if applied to these cells in mono-culture. However, this drug was rendered ineffective when HCC-1143 cells were co-cultured with the fibroblast cell, HCPF, resulting in only a 20 – 40% decrease in cell viability (orange dots in Figure 2E). A flow cytometry based analysis of cell death recapitulated this drug desensitization phenotype (Figure 2F and G). Additionally, our coculture screen identified instances in which the efficacy of etoposide is improved in co-culture conditions. For example, etoposide was ineffective in killing Mesenchymal-Like Hs578T cells in mono-culture, but killed more than 50% of these cells grown in co-culture with skin fibroblast cells, WS1 (purple dots in Figure 2E). This phenotype was interesting because our prior studies have found that etoposide, a Topo II inhibitor, is minimally active in mono-culture, which was surprising given the clinical utility of this compound (Lee et al., 2012). Flow cytometry based analysis of cell death confirmed that etoposide induced cell death in Hs578T is significantly enhanced by co-culture with WS1 fibroblast (Figure H-I).

### Principal Component Analysis highlights TNBC-fibroblast interactions as critical determinants of drug sensitivity

Prior studies that have interrogated fibroblast-tumor cell-drug interactions have found that these interactions generally result in drug resistance, with rare instances in which stromal cell interactions lead to drug sensitization (Mcmillin et al., 2010; Straussman et al., 2012). Mechanisms that contribute to this directional variability have not been identified, likely because few drug sensitizing phenotypes had been previously found. In contrast, our screen reveals that fibroblasts sensitize and de-sensitize TNBC drug response with similar frequencies (Supplemental Figure 4B). Thus, we reasoned that a statistical analysis of our data could reveal which influences account for the observed directional variation. We sought to determine if certain TNBC cells, fibroblast cells, or drugs were intrinsically more likely to be involved in sensitizing or de-sensitizing interactions (Supplemental Figures 5-7). Indeed, we found that some TNBC cells (e.g. MDA-MB-468), drugs (e.g. sunitinib), or fibroblasts (e.g. WS1) appear to be involved in directionally biased interactions. However, we wanted to integrate these insights to determine the relative importance of each of these features in our dataset.

We performed principal component analysis (PCA) on our screening data. PCA uses the correlation structure of the data to reduce data dimensionality to a smaller number of “principal components” which maximally capture the variance of the dataset (Janes and Yaffe, 2006). PCA can be used to generate a simplified description of the observed data, and here, we were particularly interested in using PCA to quantify the relative contribution of each measured influence (e.g. specific tumor cells, fibroblasts, drugs, or unique combinations of each) to the overall observed pattern of data.

PCA identified 10 principal components, with the first two components capturing 53% of the overall variation in the data. The projection of our data onto PC1 and PC2 revealed a clear separation of BL and ML cells, revealing that TNBC subtype dependent responses account for a significant portion of the observed pattern (Figure 3A and C). Notably, this expected pattern was not visible in drug response data collected on these TNBC cells grown in mono-culture (Figure 1B and 3B-C). Thus, PCA suggest that differences in chemosensitivity that are commonly observed between BL and ML subtypes of TNBC are not a cell intrinsic property, but rather a product of interactions between TNBC cells and stromal components such as fibroblasts.

**Figure 3:**
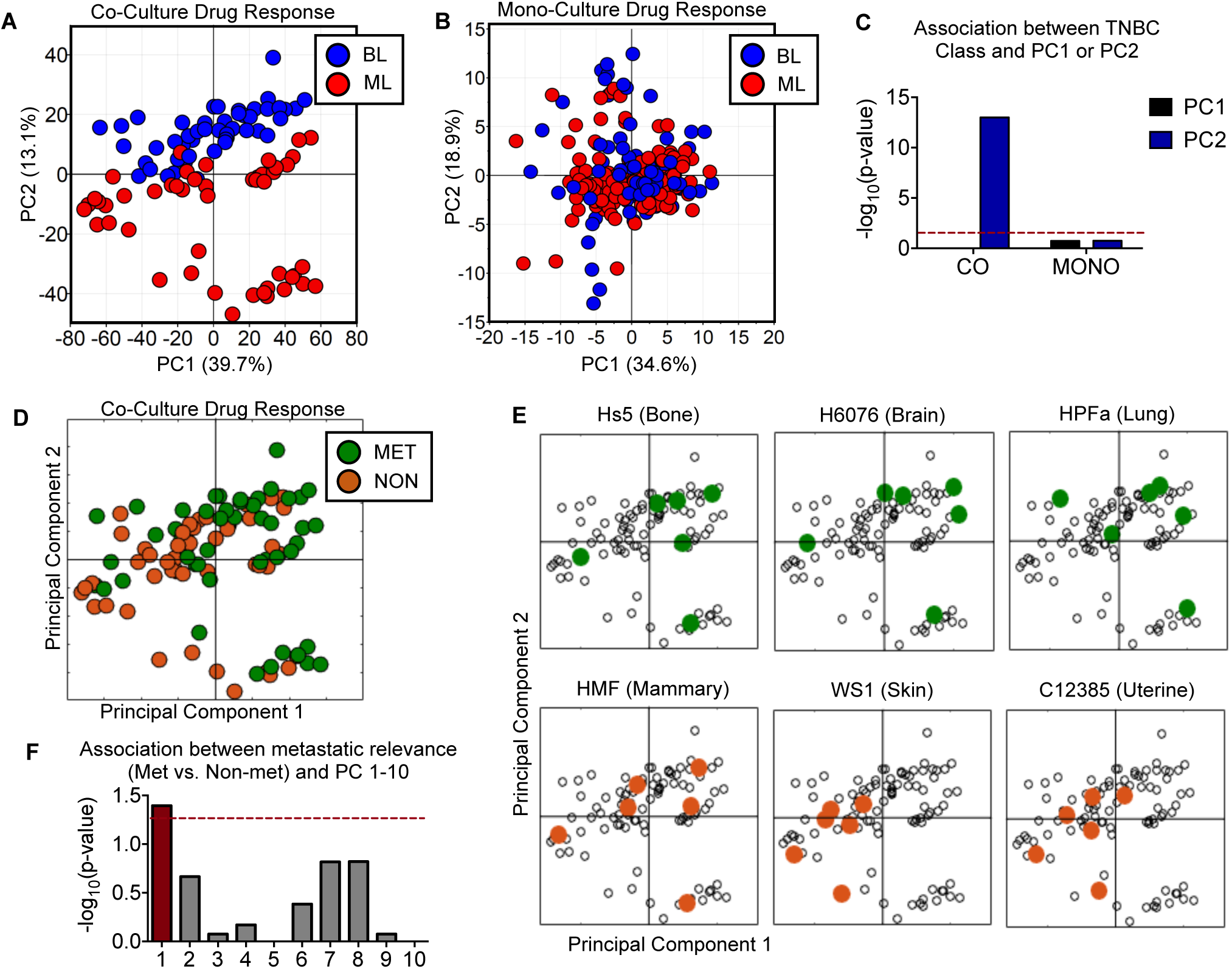
Drug responses from TNBCs treated in co-culture, but not mono-culture, accurately distinguish cells Basal-Like and Mesenchymal-Like TNBC subclasses. **(A)** PCA on co-culture drug response. **(B)** PCA on mono-culture drug response. **(C)** Association between identified principal components and Basal A subclass. p-value calculated using Fisher’s Exact test. Red line marks 0.05 cut-off. **(D)** PCA scores projection, as in panel A. **(E)** Scores for six fibroblast cell lines highlighted. Total data are shown in Supplemental Figure 8. Common metastatic locations are highlighted in green, and uncommon sites are orange. **(F)** Positive scores on PC1 are associated with common metastatic sites and negative scores on PC1 are associated with uncommon sites (p = 0.040). P-values generated using Fisher’s Exact Test.

Another important insight derived from PCA of TNBCs grown in co-culture was that the variation associated with BL vs. ML subtypes was captured exclusively on PC2, rather than PC1 (Figure 3C). By definition, PC1 captures information that is unrelated to PC2, and in this dataset, PC1 captures a three fold greater amount of the data variance compared to PC2 (39% of total dataset versus 13%). Statistical enrichment analysis revealed that PC1 captures variation associated with each of the 16 primary fibroblast cells, generally separating fibroblasts derived from common metastatic sites from those derived from organs not typically associated with breast cancer metastases (Figure 3D-F, and Supplemental Figure 8B-C). Subsequent principal components – PC3-10, which collectively account for 25% of the data – were not associated with the dichotomy of “Met vs. Non-Met” stroma, or “BL vs. ML” TNBCs, but instead were associated with specific tumor cell line-fibroblast-drug interactions. Taken together, PCA analysis reveals that, while gene expression states and anatomical location both play strong roles in modulating drug response, influences induced by fibroblasts account for a greater portion of the response variability than influences associated with the intrinsic differences between the Basal-like and Mesenchymal-Like gene expression of tumor cells.

### Anatomical origin of fibroblast dictates the directionality of drug influence

PCA revealed that differences in drug sensitivity were strongly associated with the anatomical origin of fibroblast cells, but from this analysis it was not clear what aspect of drug sensitivity was associated with fibroblast origin. In other words, from the PCA data it is not clear if fibroblasts alter the magnitude of TNBC drug response (correlated X-Y variance), or alternatively, the degree to which TNBC drug response was altered by co-culture (uncorrelated X-Y variance). To inspect this further, we arrayed all data clustering each unique cancer-fibroblast-drug combination by dose and time, in order to highlight conserved fibroblast-dependent influences (Figure 4A). Each data tile was then subsequently grouped by stromal location and drug, and a map was created for each TNBC cell line to facilitate visual inspection of the relative influences induced by each fibroblast line. To test whether fibroblast origin was associated with differences in the magnitude of drug response, we generated maps using the percent viability in co-culture (i.e. y-axis data from Figure 2E). From these maps, differences between fibroblast lines were not apparent, suggesting that fibroblast origin does not alter drug sensitivity magnitude (Supplemental Table 3). Next, to test if fibroblast origin was associated with the degree to which drug responses were altered in co-culture, we generated maps using the co-culture:mono-culture response ratio (Figure 4B and Supplemental Figure 9). From this analysis, a clear difference between fibroblast lines was visible, with fibroblasts derived from common sites of metastasis generally enhancing survival of TNBCs, whereas those derived from other organs generally were neutral or enhanced TNBC cell death. Interestingly, these patterns were observed across nearly all drugs, revealing that the location specific trends were more robust than drug specific responses. To determine if these visual trends were statistically robust, we calculated the mean response ratio for each unique cancer-fibroblast-drug combination, and separated these data by metastatic location (Figure 4C). These data confirmed a statistically significant difference in the directionality of influence between fibroblasts derived from organs that are common or uncommon metastatic sites. Additionally, we also performed statistical analyses including only the 5039 drug responses, which were the largest and most significantly changed phenotypes (Supplemental Figure 4D). This analysis of extreme “outliers” revealed that, while the total number of “hits” were similar between metastatic and non-metastatic sites, the directionality of these hits was significantly different between these groups, and consistent with the insights generated using average response ratio (Figure 4D). Taken together, these results are consistent with those from PCA, and further reveal that fibroblasts induced fundamentally opposing influences on the drug response of TNBC cells, largely dependent on the anatomical origin from which the cells were harvested.

**Figure 4:**
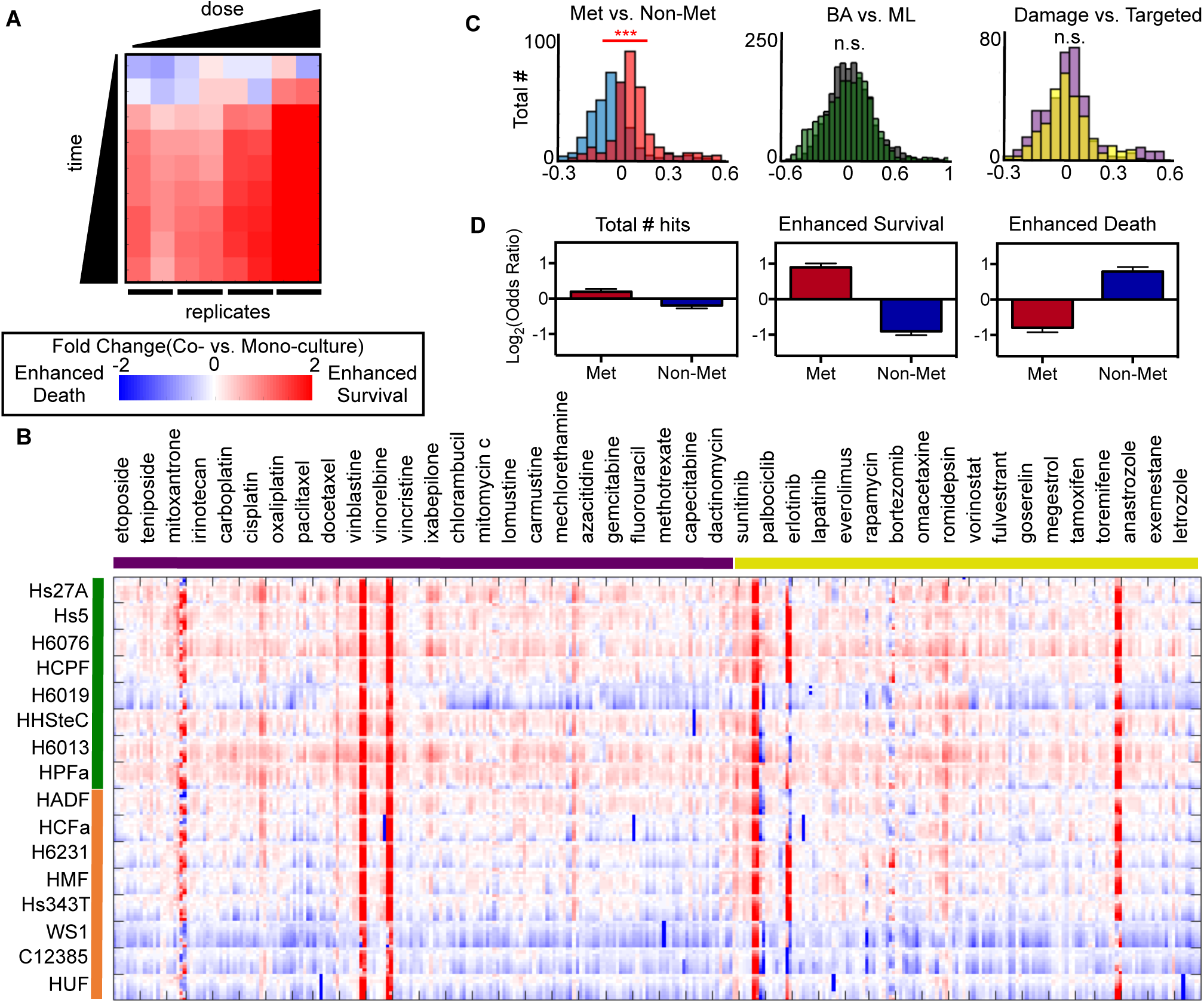
Divergent interactions between TNBCs and fibroblasts from common vs. uncommon metastatic locations (A and B) Ratio of co-culture vs. mono-culture drug response. 72 data points for each unique cancer-fibroblast-drug interaction arrayed by dose and time (9 time points and 4 doses in duplicate). **(A)** Example of total data for BT20-Hs27a-cisplatin. **(B)** Average across all cell lines. See Supplemental Figure 9 for additional cell line specific data. Top 8 fibroblasts (green bar, left) are from common metastatic organs; bottom 8 fibroblasts (orange bar, left) are from uncommon metastatic sites. 24 left-most drugs (purple bar, top) are cytotoxic chemotherapies; 18 right-most drugs (yellow bar, top) are targeted therapies. **(C)** Distributions of fibroblast influence for metastatic vs. non-metastatic locations (Met vs. Non-met), Basal-like versus Mesenchymal-Like (BL vs. ML), and cytotoxic versus targeted therapies (Damage vs. Targeted). Data are the mean co-culture:mono-culture survival ratio for each unique cancer-fibroblast-drug interaction. *** p <0.001; n.s. = not significant. **(D)** Degree of enrichment for fibroblasts from Met vs. Non-met locations within 5039 significantly altered drug responses (“Total # Hits”; See Supplemental Figure 4D), or from the subset of these that increased survival or increased death. Data shown are Odds Ratios from Fisher’s Exact Tests.

### Fibroblasts alter TNBC drug response through modulation of the mitochondrial priming state of TNBC cells

Next, we aimed to determine the mechanism by which fibroblasts interact with TNBC cells to produce divergent and largely drug-independent modulation of drug response. The simplest mechanism that is consistent with our observations would be that these fibroblasts cause a direct TNBC cell growth or survival defect, independent of the drugs added (Figure 5A, example i). To test this, we used GFP-tagged TNBC cells to monitor TNBC specific growth/survival phenotypes. We found that most fibroblasts cells either did not alter the growth rate of TNBC cells or induced a modest growth rate increase of TNBC cells grown in co-culture (Figure 5B). Furthermore, for fibroblasts that consistently sensitized drug response rates in all TNBC cell lines (WS1, C12385, or HUF), co-culture did not significantly alter growth or survival, suggesting that a fitness or survival defect does not account for the broad-spectrum drug sensitization seen in co-culture with these cells. In rare instances co-culture conditions did result in a significant TNBC cell growth rate decrease, such as seen with MDA-MB-231 cells grown with H6013, a fibroblast derived from lung tissue (Figure 5B). Notably, the M231-H6013 interaction induced broad-spectrum drug desensitization (i.e. enhanced survival, see Supplemental Figure 9). Thus, even in the rare instances in which fibroblast cells mediated fitness defects, growth rate or survival modulation does not appear to account for the observed pattern of influences on TNBC drug response.

**Figure 5:**
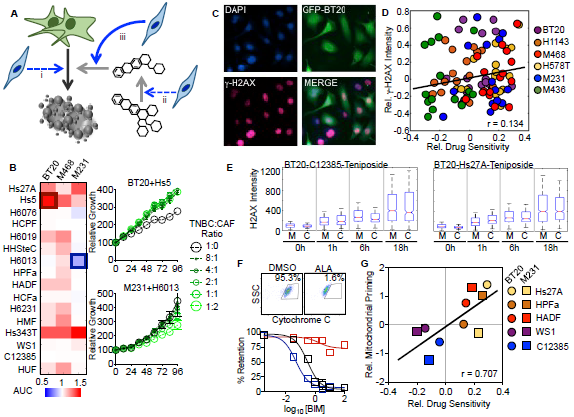
Fibroblasts alter drug sensitivity through modulation of mitochondrial apoptotic priming. **(A)** Schematic of possible mechanisms by which stromal cells alter drug sensitivity. **(B)** TNBC cells labeled with GFP were grown in mono-culture or in co-culture with listed fibroblasts. Growth rate quantified using well fluorescence. Heatmap data are area under curve (AUC) from biological triplicate measurements. Cells plated at 1:1 ratio. Growth curves shown for the most enhanced and suppressed growth rate. **(C-E)** γH2AX (p-H2AX, S139) monitored by immunofluorescence microscopy. **(C)** Representative image of GFP-BT20 cells co-cultured with HADF. **(D)** Quantification of TNBC nuclear γH2AX, using automated image analysis (CellProfiler). Scatterplot of co-culture:mono-culture viability ratio (from screen in Fig. 2E) compared to coculture:mono-culture H2AX intensity ratio. Both plotted in Log2 scale. Average number of nuclei per counted per condition is 758 (range 93 – 1632) **(E)** Boxplots showing distribution of nuclear γH2AX intensities over time for BT20 cells in mono-culture (M) or in co-culture (C) with strongly drug sensitizing fibroblasts (C12385) and de-sensitizing fibroblasts (Hs27A). **(F)** Mitochondrial priming assay. (top) Cytochrome C retention quantified using the iBH3 profiling assay. Alamethicin (ALA) used as a positive control for mitochondrial rupture. (bottom) Mitochondrial priming quantified using exposure to varied concentrations of BIM. **(G)** Scatterplot comparing relative drug sensitivity (as in panel E) compared to degree of co-culture induced change in mitochondrial priming. Priming status quantified as AUC from BIM dose response. Data are from biological quadruplicates.

A second mechanism by which fibroblast cells could enhance drug efficacy could be by metabolizing the drugs, creating a more potent or more bioavailable compound (Figure 5A, example ii). This mechanism was recently reported to explain a microbiome-drug interaction that modulates toxicity of the chemotherapeutic 5-FU (García-González et al., 2017). To test the role of fibroblast-mediated drug metabolism, we focused on drugs that activate death through induction of DNA damage. For this set of compounds, drug potency should be proportional to level of γ-H2AX, which marks sites of DNA double stranded breaks. We quantified γ-H2AX nuclear intensity in the presence and absence of fibroblast co-culture. These measurements were made at 4 time points following exposure to teniposide, a Topo II inhibitor similar to doxorubicin, which is used clinically in the treatment of TNBC (doxorubicin fluorescence limits use of this compound). We used GFP-labeled TNBC cells to identify TNBC cell nuclei and images were quantified using automated image analysis (CellProfiler, (Lamprecht et al., 2007)). Overall, we found many cases where fibroblasts modulated γ-H2AX levels (Figure 5D). Importantly, however, the degree to which γ-H2AX was modulated by fibroblast cells was poorly correlated with the phenotypic influence of these fibroblasts (Figure 5D). Furthermore, we inspected the most strongly sensitizing and de-sensitizing co-culture environments to determine if these extreme cases could be explained by differences in the apparent drug potency. TNBC nuclear γ-H2AX intensity was similar in BT-20 cells co-cultured with C12385 and Hs27A, fibroblasts that strongly sensitized and desensitized drug responses, respectively. Thus, it does not appear that fibroblast influences on drug sensitivity generally occur through modulation of the drugs themselves.

The insights gained from γ-H2AX intensity are also consistent with our general observation that fibroblast cells influence drug sensitivity in similar ways across diverse classes of drugs. In other words, it does not appear that the mechanisms by which fibroblast cells influence the drug responses in TNBC cells are specific to the drug compounds themselves or drug-specific responses of TNBC cells. Drug-induced cell death is the product of at least two independent influences: the drug-specific cell response (i.e. the ability of a drug to change a cell from a healthy to a dead state) and the degree to which the cell is “primed” for death (i.e. how “close” the healthy cell is to dying) (Chonghaile et al., 2011; Montero et al., 2015). Thus, a third mechanism that we tested was whether fibroblasts alter the degree of mitochondrial apoptotic priming. We used the BH3 profiling technique to measure changes in the relative state of mitochondrial priming (Ryan and Letai, 2013).

This assay quantifies the amount of recombinant BH3, a pro-apoptotic peptide, required to rupture mitochondria. We selected five fibroblast cells that produced the strongest and most consistent modulation of drug sensitivity. The mitochondrial response to BIM peptide was quantified by monitoring cytochrome c retention by flow cytometry (Figure 5F). BH3 profiling revealed that fibroblast co-culture conditions significantly altered the mitochondrial priming state of BT-20 and MDA-MB-231 cells. Furthermore, the degree to which mitochondrial priming was increased or decreased was also highly correlated with relative drug sensitivity observed in our co-culture screen (Figure 5G).

### Fibroblast-secreted IL8 induces hyper-sensitivity to DNA damaging chemotherapy

WS1 (skin), C12385 (uterine), and HUF (uterine) cells, sensitized all six TNBC cell lines to nearly every drug tested. We prioritized understanding mechanisms by which these cells induce broad-spectrum drug sensitization, as these may be therapeutically relevant insights. To begin, we determined whether conditioned media from these fibroblast cells also induced drug sensitization. Conditioned media was collected following 48 hours of culture with fibroblasts and added to TNBC cells prior to addition of teniposide. To measure the rate of TNBC cell death, we used Sytox green, a cell impermeant dye that is fluorescent only when bound to DNA. We found that conditioned media from WS1, HUF, and C12385 sensitized all six TNBC cell lines tested to teniposide (Figure 6A). In contrast, conditioned media from Hs27A, a bone fibroblast cell, generally did not alter TNBC drug response, although these data were more variable across TNBC cells.

**Figure 6:**
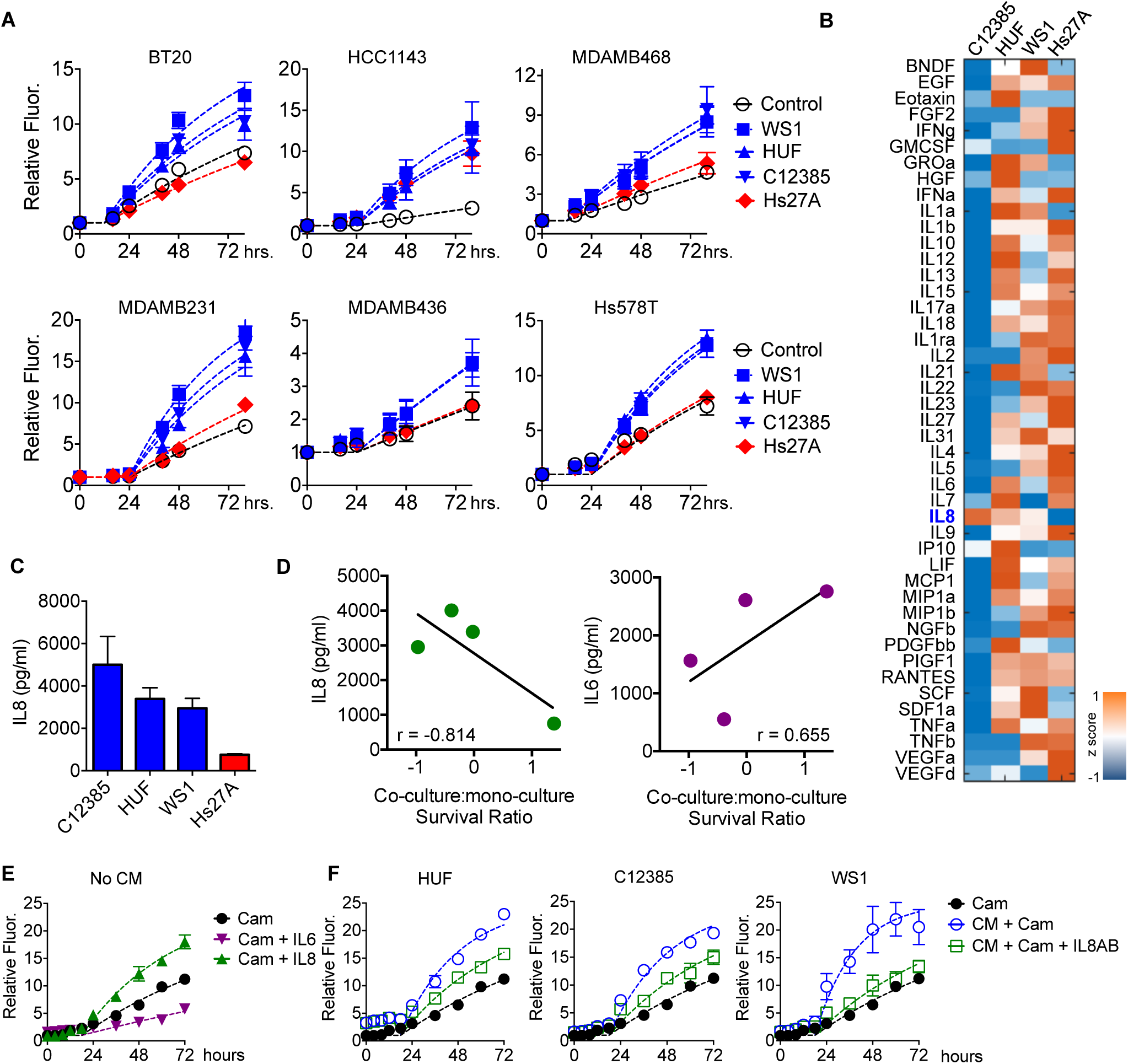
IL-8 secretion from fibroblast cells enhances sensitivity to DNA damage. **(A)** Cell death measured using Sytox green following exposure to 10 µM teniposide, in the presence or absence of fibroblast conditioned media (CM). **(B and C)** Cytokine and growth factor secretion quantified using Luminex assay. Heatmap in panel B is colored according to the relative secretion across four CM samples. Data are from biological duplicate samples. **(D)** Secretion of IL8 and IL6 are shown relative to the Log2 survival ratio (data are from co-culture drug screen as in Figure 2E). **(E and F)** Cell death measured using Sytox green following exposure to 5 µM camptothecin (Cam), in the presence or absence of recombinant IL6 or IL8 (E), or conditioned media (CM) and/or IL8 neutralizing antibody (IL8AB). IL6 and IL8 were used at 50 ng/µL, and IL8AB was used at 100 µg/ mL.

To identify secreted factors that are responsible for these phenotypes, we profiled conditioned media for the presence of 45 common cytokine, chemokine, and growth factors (Figure 6B). We reasoned that the relative secretion profile for factors that induce drug sensitization should be inversely correlated with the relative drug sensitivity observed in our co-culture screen. Of the 45 cytokines profiled, we observed strong negative correlation only for IL8 (Figure 6C and D). Consistent with this line of reasoning, we also observed a positive correlation between relative drug sensitivity and secretion of IL6, a cytokine that is already known to induce resistance to DNA damaging chemotherapies (Figure 6D) (Gilbert and Hemann, 2010). To test if IL8 also alters sensitivity to DNA damage, we again used Sytox green to quantify the rate of cell death. Recombinant IL8 increased the rate of camptothecin induced cell death by approximately 2.5-fold, whereas recombinant IL6 decreased the rate of cell death by approximately 2-fold (Figure 6E). To further determine the role of IL8 secretion in the drug sensitization phenotype observed in conditioned media, we tested IL8 neutralizing antibodies with conditioned media from WS1, HUF, or C12385 fibroblasts. In all three cases, IL8 neutralizing antibodies significantly inhibited the drug sensitization induced by conditioned media (Figure 6F). Notably, IL8 neutralizing antibodies failed to restore drug sensitivity to the levels observed in the absence of fibroblast conditioned media, suggesting that other secreted factors also contribute to the drug sensitization phenotype.

## DISCUSSION

In this study, we explored interactions between tumor cells and stromal cells to identify those that modulate sensitivity to commonly used chemotherapeutics. We found that fibroblasts can alter drug sensitivity of tumor cells, and that the responses were highly variable, both in magnitude and in direction. Our statistical analysis clarified that the directional variability in fibroblast influence is predominantly associated with the anatomical organ from which the fibroblast cells were harvested. Specifically, fibroblasts from common sites of metastasis typically desensitize tumor cell drug responses. This interaction was fundamentally different than what was observed with fibroblasts from organs that do not typically accommodate metastatic growth, which typically sensitized tumor cell drug responses. Somewhat surprisingly, these influences were consistently observed, regardless of which drug was applied, which was caused by fibroblast-dependent modulation of mitochondrial priming within cancer cells. Our analysis of a small set of common cytokines shows that IL8 and IL6 contribute to drug sensitization and de-sensitization, respectively.

The contribution of cancer associated fibroblasts to a variety of tumor phenotypes has been studied, including using high-throughput co-culture based systems (Mcmillin et al., 2010; Straussman et al., 2012). Our data, in many respects, reveal phenotypes that are similar to what has been uncovered in prior high-throughput co-culture based screens. A notable exception is that, compared to prior studies, our screen revealed a greater proportion of drug sensitizing interactions between stromal and cancer cells. This difference could have resulted from the depth of our screen, as strong drug sensitizing phenotypes are also rare in our data. Alternatively, it is also possible fibroblast mediated drug sensitization is a more common phenotype in TNBC cells, as this cancer subtype was not deeply profiled in prior studies. Another possibility is that our screening methodology, which was designed to exclusively monitor drug induced cell death, contributed to the enhanced resolution of cell death sensitization. In fact, this feature may be likely to play a part given the limited ability of other approaches to quantify differences in degree of cell death. One important point regarding our assay is that the use of JC-1 to monitor environmental influences on cell death left our assay incapable of quantifying other important changes, including alterations to proliferation rate. Thus, future attempts to study tumor-stroma interactions may benefit from the use of multiple complementary screening approaches.

Nonetheless, the findings from our study have important implications that should be considered in the context of “personalized” or “precision” medicine. These concepts are explored typically using genomic analyses of tumor cells. Our data suggest that interactions between tumor and stromal cells often alter drug sensitivity, and moreover that these interactions are a potentially more potent source of variation in drug sensitivity than tumor cell gene expression state. Thus, our data suggest that personalized treatment regimens will ultimately need to consider micro-environmental features of tumors – and in particular the behaviors of stromal fibroblasts – in addition to genomic considerations. Furthermore, because fibroblasts appear to have the capacity for both positive and negative influence in modulating tumor cell drug sensitivity, simple analysis of their presence or absence is not likely to be robustly informative. Our study identifies inflammatory cytokines such as IL8 and IL6 as key modulators of tumor cell drug sensitivity. Because of the limited nature of our cytokine screen, it is extremely unlikely that these cytokines explain all, or even most, of the phenotypes identified in this study. Future studies should focus on identifying a more comprehensive list of cell non-autonomous features that modulate drug responses in cancer cells. These insights may be valuable strategies for enhancing drug induced cell death in cancer treatment.

## MATERIALS AND METHODS

### Cell lines and reagents

Cell lines BT-20, HCC-1143, Hs-578T, MDA-MB-231, MDA-MB-436, MDA-MB-468, HCC-2157, HCC-1806, HCC-1395, Hs27A, HS-5, WI-38, IMR-90, Hs 343.T, WS-1 were obtained from American Type Culture Collection (ATCC) and cell line CAL-120 was obtained from Deutsche Sammlung von Mikroorganismen und Zellkulturen GmbH (DSMZ). All cell lines were grown in 10% FBS (Thermofisher Hyclone cat# SH30910.03 lot# AYG161519), 2 mM glutamine, and penicillin/streptomycin. BT-20, CAL-120, and WS-1 were cultured in Mem*α* + Earle’s Salts. HCC- 1143, HCC-2157, HCC-1806, HCC-1395 were cultured in RPMI 1640 media Hs-578T, MDA-MB-231, MDA-MB-436, MDA-MB-468, Hs27A, HS-5, Hs 343.T were cultured in Dulbecco’s modified eagles medium (DMEM). Hs578T were further supplemented with 10ug/ml insulin. Primary fibroblasts, H-6231, H-6201, H-6076, H-6019, and H-6013 were purchased from Cellbiologics; HCPF, HPF-a, HHSteC, HMF, HAdF, HUF, and HCF-a were purchased from ScienCell; and C- 12385 was purchased from Promocell. Primary fibroblast cells purchased from Cellbiologics, ScienCell and Promocell were cultured in the media (Sciencell - Fibroblast Medium cat# 2301; Cellbiologics - Complete Fibroblast Medium /w Kit cat# M2267; Promocell - Fibroblast Growth Medium 2 cat# C-23020) for 4 doublings before being transitioned to DMEM. All cells were cultured at 37C in a humidified incubator supplied with 5% CO2 and maintained at a low passage number (less than 20 passages for cancer). Prior to expansion and freezing, a small sample of each primary fibroblast was expanded to determine each cell’s Hayflick limit to ensure that experiments could be performed prior to the onset of replicative senescence. A complete list of drugs used in this study is included in Supplemental Table 2; antibodies and other reagents used in this study are listed in Supplemental Table 5.

### Co-culture screen using JC1 dye

Fibroblast cell lines were grown to 80% confluence before being trypsinized, and stained with 5µM CellTrace Violet Proliferation dye (Thermofisher #C34557) in PBS at a concentration of 1x10^6 cell/mL for 15 minutes at 37C. 1500 stained cells were plated in 40µL FluoroBrite media (Thermofisher # A1896701), supplemented with 10% FBS, 2mM glutamine and penicillin/streptomycin, in a Greiner clear 384 well plate (#781986) and allowed to adhere for 3 hours. Cancer cell lines were then trypsinized, and stained with 1.5µg/mL (final concentration) JC-1 (Thermofisher # T3168) in FluoroBrite at a concentration of 1x10^6 cell/mL for 20 minutes at 37C. Cancer cells were then plated at 1500 cells in 40uL FluoroBrite per well in the 384 well plate. For mono-culture conditions, unlabeled cancer cells were added to each well, in order to keep the cell density consistent with co-culture conditions. Cells were allowed to adhere overnight. The following morning, 8µL of a 10x drug stock was added to the wells using a VIAFLO 96 Electronic 96-channel pipette machine. JC-1 fluorescence was then read at 5 spots across each well using a Tecan M1000 Plate Reader at the excitation wavelength of 535nM +/-17nM and an emission wavelength of 590nM +/- 17nM every 8 hours for 72 hours. Background fluorescence was determined by treating labeled cells with Alamethicin, a membrane permeabilizing agent that punctures plasma membrane and mitochondrial membranes. Fluorescence measurements were normalized relative to pre-drug treatment values for each well.

### Cell viability and cell death assays

Cell viability assays were performed either using CellTiter-Glo (cat# G7570), for cells grown in mono-culture, or flow cytometry, for co-culture assays (other than the co-culture screen, described above). For CellTiter-Glo, which measures viability as a function of ATP concentration, cells were plated in Greiner 96 well plates (cat# 655 090) at 5000 cells per well in 100uL of their respective growth media and allowed to adhere overnight. 10uL of a 10x drug stock, diluted in PBS, was added to each well. Cells were subsequently allowed to grow at 37C for 72 hours. At 72 hours post drug addition, 33uL of CellTiter-Glo reagent was added to each well. The CellTiter-Glo assay was performed according to manufacturer’s directions, with the reagent diluted 1:3 (relative to media volume). Luminescence was read using a Tecan M1000 Plate Reader. Cell death measurements to validate the JC-1 screen data were collected using the Live/Dead Violet reagent (Thermofisher cat# L34963) and analyzed by flow cytometry. Cancer cells and fibroblast cells were plated at a 1:1 ratio in DMEM and allowed to adhere overnight. Drugs were added from a 1000x stock and cells were exposed for the specified times. Cells were trypsinized at the specified times, suspended in PBS at a concentration of 1x10^6 cells per mL and stained with a 1:1000 dilution of the Live/Dead Violet reagent for 30 minutes on ice. Cells were then fixed with 4% formaldehyde for 10 minutes at room temperature and run on an LSR II FACS machine with a laser excitation of 405nm and emission of 450nm.

### Growth rate measurements using GFP labeled cells

To determine cell proliferation rate using a fluorescence plate reader, TNBC cells were stably transfected with GFP (pRetroQ-AcGFP1-N1). Transfected cells were selected with puromycin (BT- 20 at 1.5µg/mL, 468 at 0.5 µg/mL and 231 at 2 µg/mL). Cells were selected until a parallel non-transformed plate exposed to puromycin was completely dead. The selected population was subsequently sorted by FACS to collect cells with similar levels of GFP fluorescence. For co-culture experiments, fibroblast cell lines were plated at an 8:1, 4:1, 2:1, 1:1, and 1:2 ratio to cancer cells in a Greiner 96 well plate in 100uL of FluoroBrite media and allowed to adhere for 3 hours. Following adherence of fibroblast cells, TNBC cells constitutively expressing GFP were plated at a concentration of 10,000 cells per 100uL of FluoroBrite media and allowed to adhere overnight. Cell measurements were measured every 24 hours for 96 hours using a Tecan M1000 Plate reader.

### Immunofluorescence Microscopy

For quantitative analysis of p-H2AX nuclear intensity, fibroblast cells were plated at a density of 1500 cells per 25 µL in DMEM in a 384 well plate and allowed to adhere for 3 hours. Cancer cells were stained with 5 µM CellTrace CFSE dye (Thermofisher cat# C34554) at a concentration of 1x10^6 cells per mL in PBS for 15 minutes at 37C. Labeled cells were plated at 1500 cells per 25 µL DMEM and allowed to adhere overnight. Drugs were added from a 10x stock solution in PBS and cells were exposed for 1, 6, and 18 hours before being fixed with 4% formaldehyde for 10 minutes at room temperature. Cells were washed twice in PBS, then permeabilized with 0.5% Triton X100 for 10 minutes at room temperature. Cells were washed twice with PBS; blocked in 10% goat serum (Thermofisher cat# 16210064) for one hour; stained with the p-Histone H2A.X (Ser139) antibody (Cell Signaling Technologies #9718S) in 1% goat serum in PBS overnight at 4C; stained with Alexa-647 antibody (1:250 dilution, Thermofisher A21244) in 1% goat serum in PBS for 2 hours at room temperature. Imaging was performed using an IXM-XL high throughput automated microscope. Analysis was performed using a custom CellProfiler pipeline (available upon request).

For α-SMA staining, fibroblast cells were stained with 5 µM CellTrace Far Red dye (Thermofisher cat# C34564) and cancer cells were stained with 5uM CellTrace Violet dye, each as described above. Each cell type was plated at a 1:1 ratio. Cell fixation, permeabilization, and staining were performed as above for H2AX. Cells were stained with the α-SMA antibody (Cell Signaling Technologies #19245S) in 1% goat serum in PBS overnight at 4C.

### Mitochondrial priming assays

Mitochondrial priming assays were performed according to the iBH3 protocol from Ryan et al (Ryan et al.). For monoculture conditions, 1x10^6 cancer cells were plated in a 10cm dish and allowed to adhere overnight. For co-culture conditions, fibroblasts were stained with Cell Trace Violet plated at a 1:1 ratio with cancer cells. 24 hours post plating cells were trypsinized and arrayed in a 384 well plate. A dose series of BIM peptide was added (100uM, 33uM, 10uM, 3.3uM, 1uM, 0.33uM) along with either DMSO (vehicle control) or alamethicin, a mitochondrial depolarizing agent, which was used at a final concentration of 25uM as a positive control. Plasma membrane permeabilization was achieved by the addition of digitonin at a final concentration of 20 µg/mL. Cells were incubated with BIM peptide at room temperature for 1 hour before being fixed and stained for cytochrome c retention (Fisher cat# BDB560263). Samples were analyzed on an LSRII flow cytometer.

### Conditioned media assays

Fibroblasts were plated in a 10 cm dish at a concentration of 750,000 cells in 10mL of DMEM and allowed to adhere for 3 hours. For co-culture conditions, following fibroblast adherence, 750,000 cancer cells were added to the plates. Cultures were incubated at 37C for 48 hours and conditioned media was collected and filtered through a 0.45um syringe filter. To test whether conditioned media altered drug sensitivity, cancer cell lines were plated in the respective conditioned media at a concentration of 3000 cells per 50uL conditioned media in a 384 well plate and allowed to adhere overnight. 24 hours post plating 5uL of a 10x drug stock was added to each well along with the Sytox green reagent (Fisher Scientific, S7020). Sytox Green was used at 5uM final concentration, and fluorescence was measured using a Tecan Plate reader. The Sytox Green fluorescence reading is proportional to the number of dead cells in the well. To determine numbers of live cells and the percent viability, Triton x100 was added to each well (0.2% for 3 hours at 37C) to induce permeabilization of remaining live cells.

### Cytokine analysis

Conditioned media for the selected fibroblast lines were collected 48 hours post plating and arrayed in biological replicates with two technical replicates each. Cytokine and chemokine analysis was performed according to the manufacturer’s instructions for the Cytokine/Chemokine/Growth Factor 45-Plex Human ProcartaPlex™ Panel 1 kit obtained from Thermofisher (cat# EPXR450-12171- 901).

### Data analysis and statistics

All statistical analyses were performed using GraphPad Prism and/or MATLAB, generally using pre-built functions (FisherTest, t test, etc.). PCA was performed using SIMCA and data were z scored (mean centered and unit variance scaled). Hierarchical clustering was performed using Spotfire using the default settings (UPGMA clustering method; Euclidean distance measure; Average value ordering weight; z-score calculation normalization method; empty value replacement: NA). Analysis of flow cytometry data was performed using FloJo.

## ACKNOWLEDGEMENTS

We thank members of PSB and the DFCI CCSB for valuable discussions during the design and execution of this study. Additionally, we thank the UMassMed School Flow Cytometry Core and High-Throughput Imaging Core for training and advice during the execution of this study. This project was conceived by BDL and MJL. Experiments were designed by BDL, ADS, SRP, and MJL. Co-culture screen was designed and executed by BDL. All other experiments were executed by BDL, TL, RR, PCG, and GR. Analysis was conducted by BDL, TL, RR, and MJL. Manuscript was written and edited by BDL, ADS, SRP, and MJL. This work was supported by grants to MJL form the Richard and Susan Smith Family Foundation, Newton, MA, the Breast Cancer Alliance, and the American Cancer Society (RSG-17-011-01). BDL was supported by a NIH training grant (Translational Cancer Biology Training Grant, T32-CA130807). Additional support was provided by the UMass President’s S&T Fund to MJL and SRP.

## Competing Interests

None

---

**Supplementary Figure 1:**
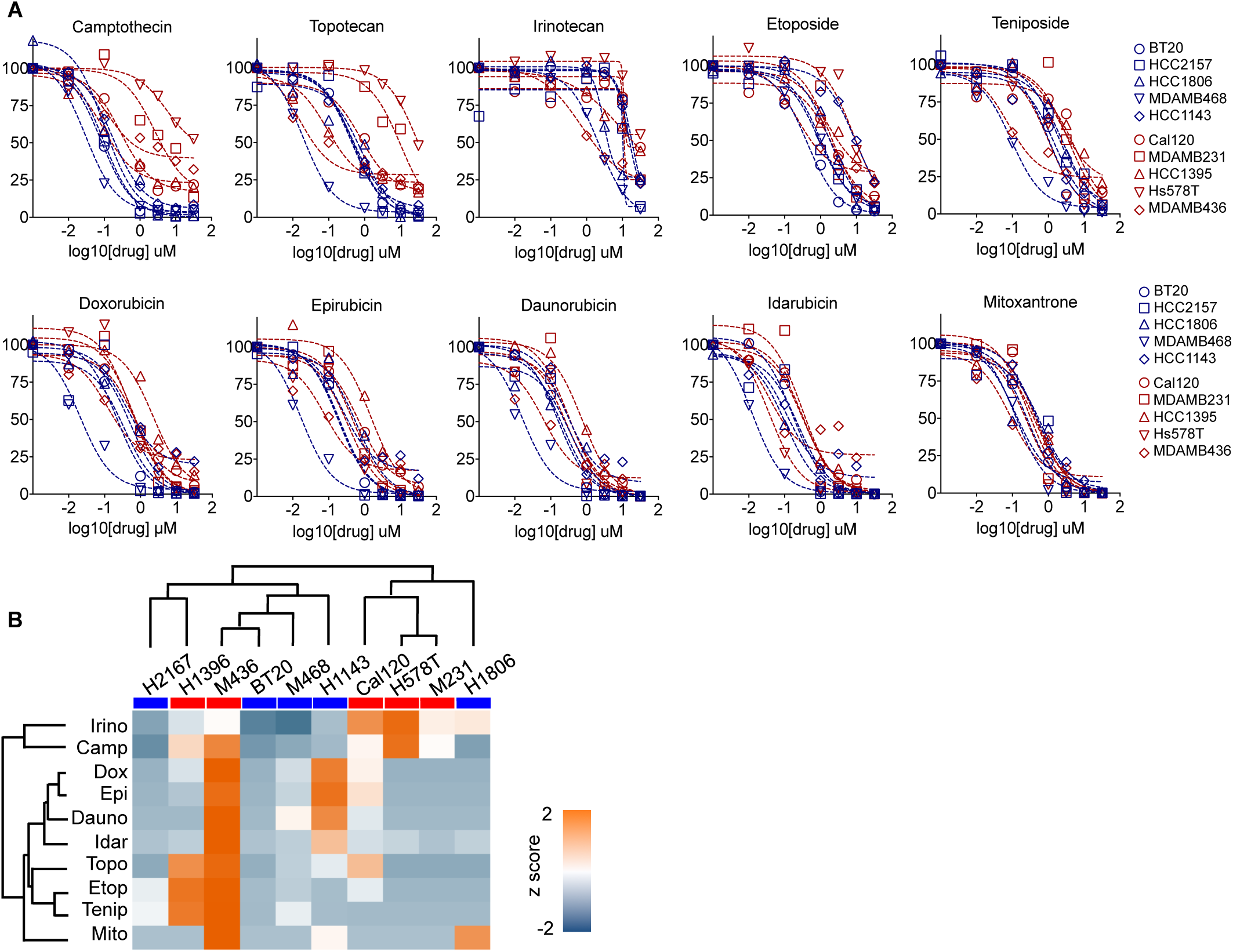
Variable sensitivity to common DNA damaging agents in TNBC cells. **(A)** Drug dose response curves for 10 Topo I and Topo II inhibitors. Data are presented as in Figure 1A. **(B)** Cell viability measured as in (A) for 10 Topo I/II inhibitors. Data are z scored max death at 72 hours (Emax). Dendograms from hierarchical clustering shown for drugs and for cells (Basal A cells highlighted with blue bar; Claudin-low cells highlighted with red bar).

**Supplementary Figure 2:**
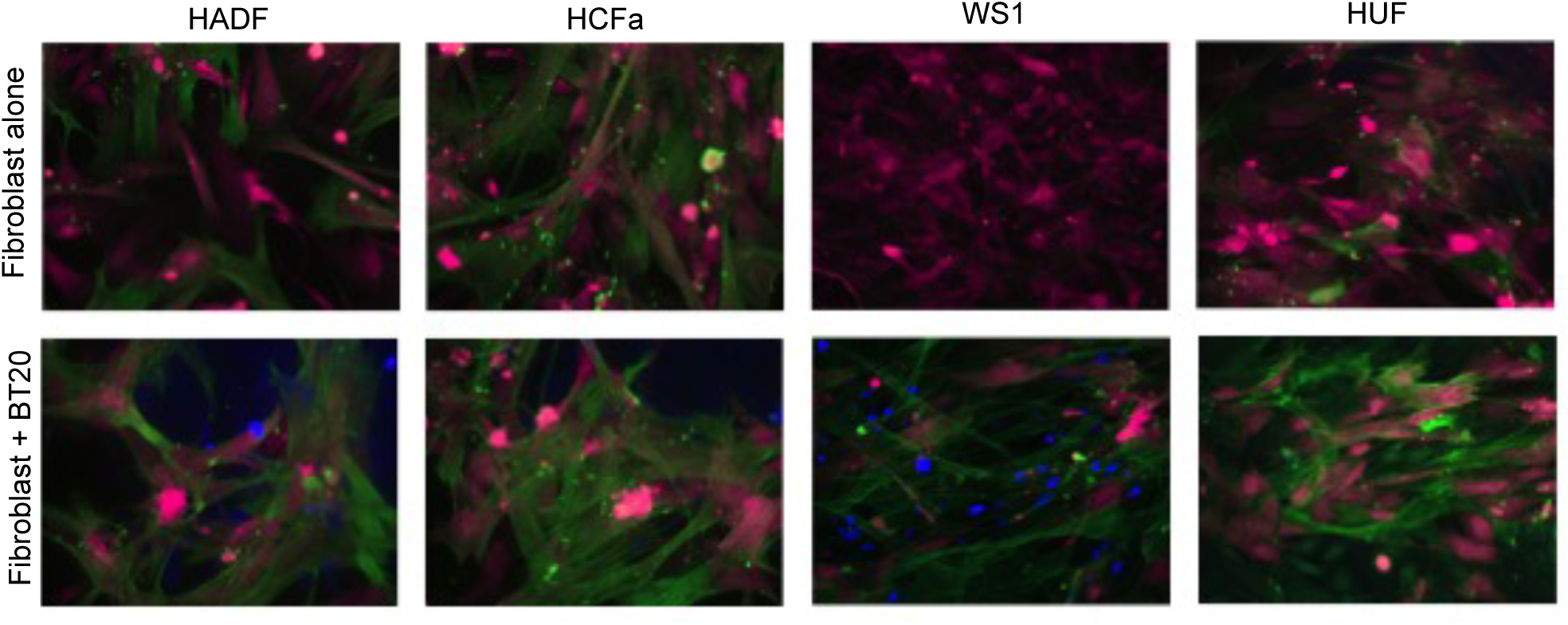
Primary fibroblasts stain positive for markers of activation in co-culture with TNBC cells. Primary fibroblasts grown in mono-culture or in co-culture with BT20 cells were stained for expression of α-smooth muscle actin (SMA), a marker for activated fibroblasts (a.k.a. “Cancer Associated Fibroblasts”, CAFs, or myofibroblasts). SMA in green. FarRed dye (CellTrace) used to stain fibroblasts and Blue dye (CellTrace) used to stain BT20. BT20 cells grown in mono-culture did not stain positive for SMA expression (data not shown).

**Supplementary Figure 3:**
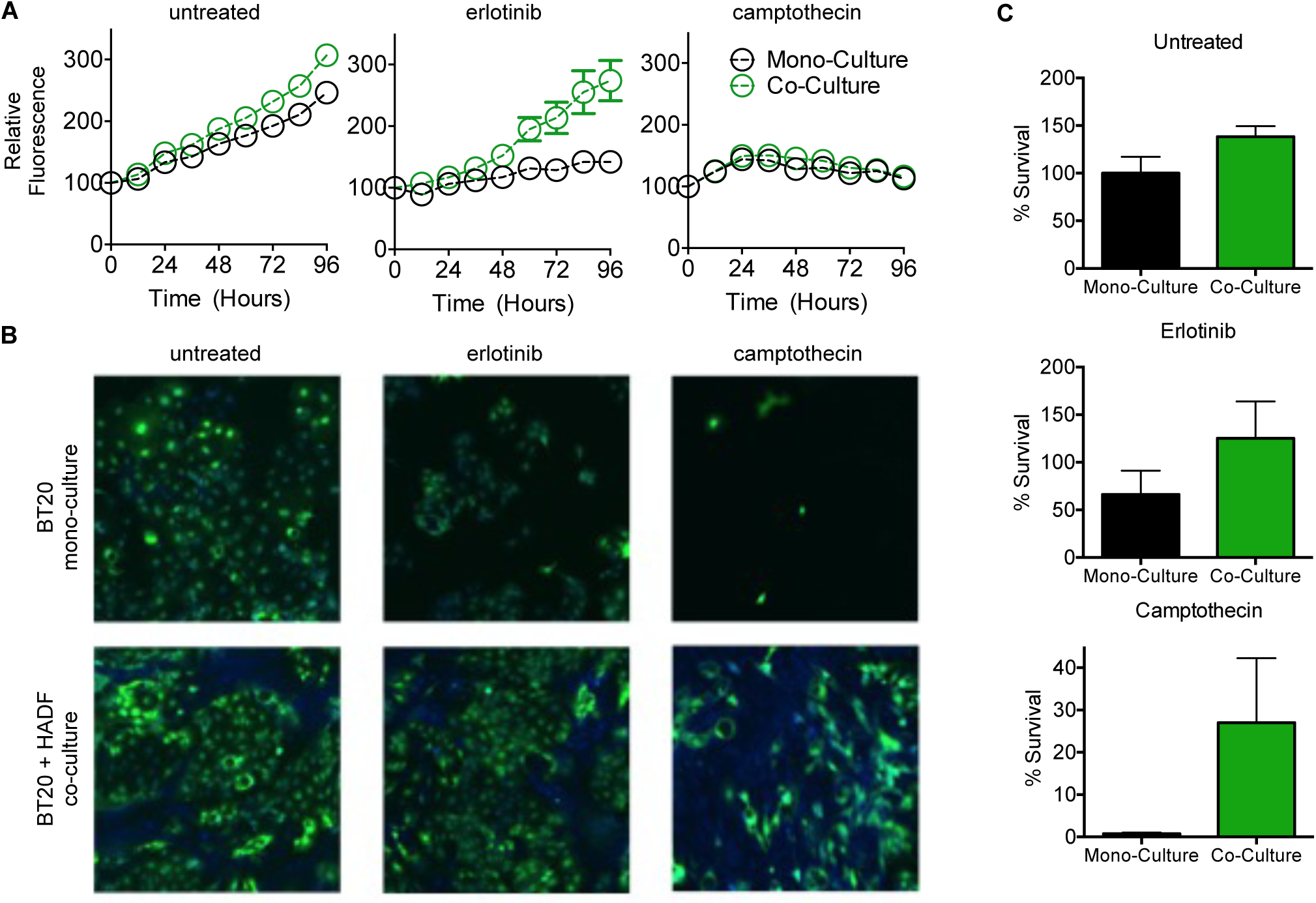
GFP-based measurements do not accurately report cell death. **(A)** Quantification of cell number using GFP fluorescence measured via fluorescence plate reader. Data are from biological quadruplicate experiments. Mono-culture is BT20 cells labeled with GFP; co-culture is GFP-BT20 + HADF, seeded at a 1:1 ratio. Data are GFP fluorescence per well in untreated cells, or following exposure to 10 µM erlotinib or 500 nM camptothecin. **(B)** Representative images from experiment in panel (A). Images collected using fluorescence microscopy following the 96 hour measurement on a plate reader. GFP-BT20 are green; HADF are stained blue using a whole cell stain. **(C)** Cell viability based on quantitative image analysis. Wells from experiment in panel (A) were imaged at 96 hours and cell numbers were quantified using CellProfiler. % Survival is relative to untreated cells grown in mono-culture. At least 300 cells were counted in every image, except BT20 mono-cultures treated with camptothecin, where the average number of cells per image was 20 (8 images collected). Well-based fluorescence measurements taken using a plate reader accurately capture proliferation phenotypes but fail to capture differences in cell death.

**Supplementary Figure 4:**
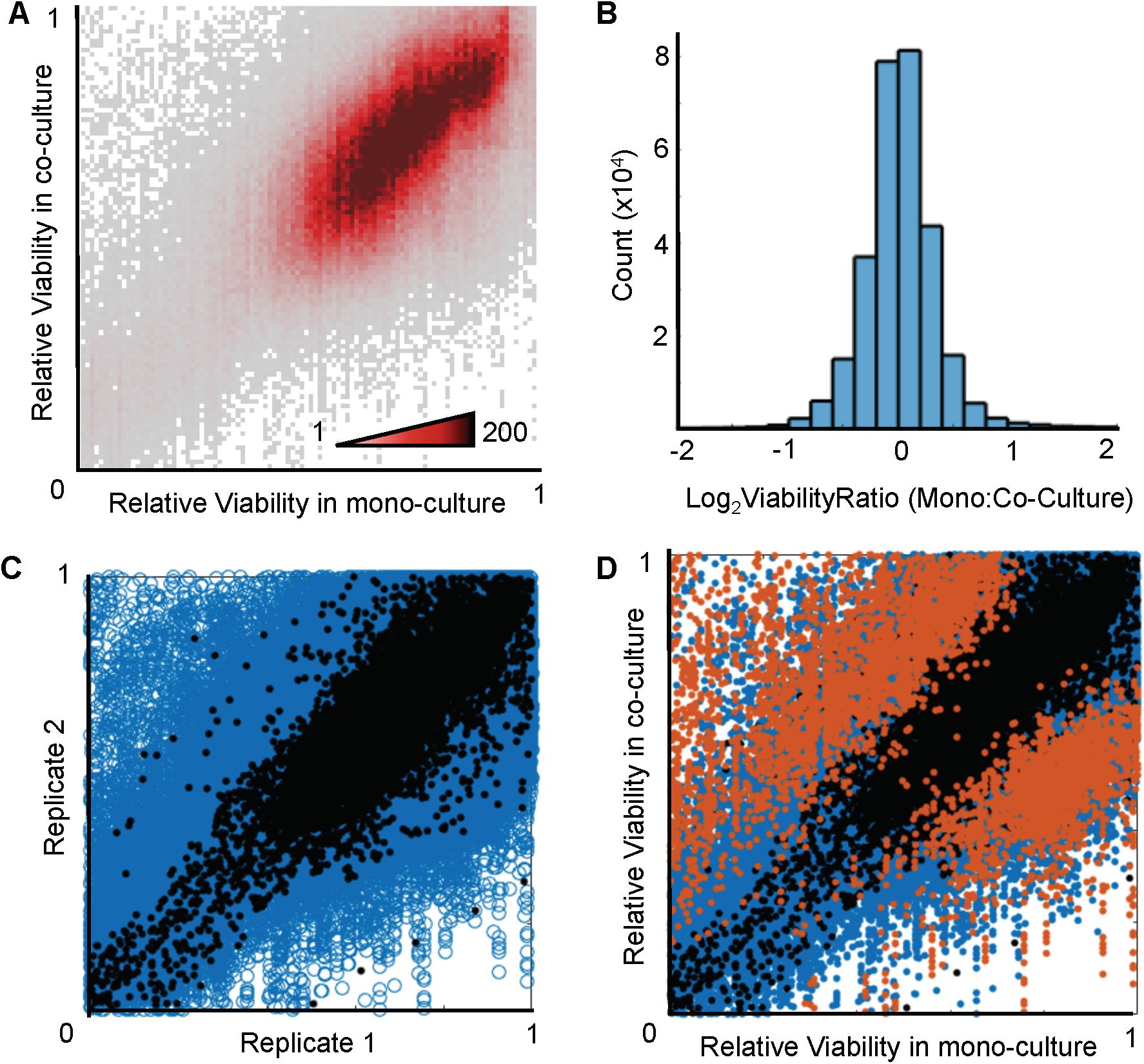
Statistical analysis of co-culture screen to identify CAF-cancer interactions that significantly alter drug sensitivity. **(A)** Density plot of co-culture screen data. 95% of 312,120 data points fall within a single dense cluster (20-50% death; no influence of CAF).Ratio of drug response in mono-culture vs. co-culture with CAFs. Data are normally distributed. Correlation among replicates (r2 = 0.7315). Two biological replicates of mono-culture drug response are shown in black (total dataset in blue, shown as reference). **(D)** CAF-cancer interactions that significantly alter drug response shown in orange (z score of mono:co-culture ratio> 3). 5039 drug responses were significantly altered relative to error among control replicates (black). See also Table XX.

**Supplementary Figure 5:**
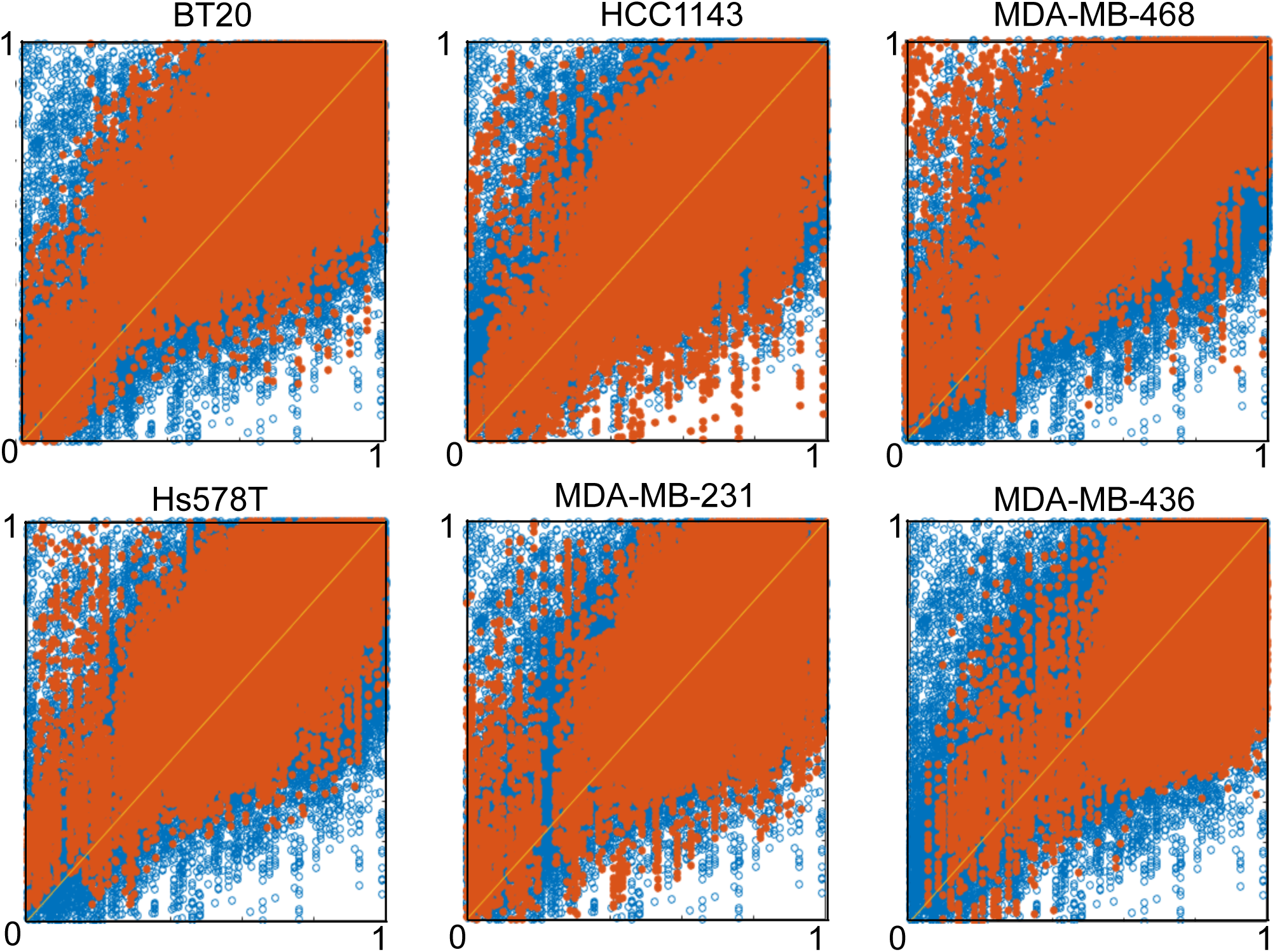
CAFs influence drug sensitivity across TNBC cell lines. Data plotted as in Figure 2E, with drug responses of TNBC cells grown in mono-culture on the x-axis, and responses in co-culture with CAFs on the y-axis. In each plot, the overall dataset is shown in blue circles and the data for each TNBC cell line is shown in orange.

**Supplementary Figure 6:**
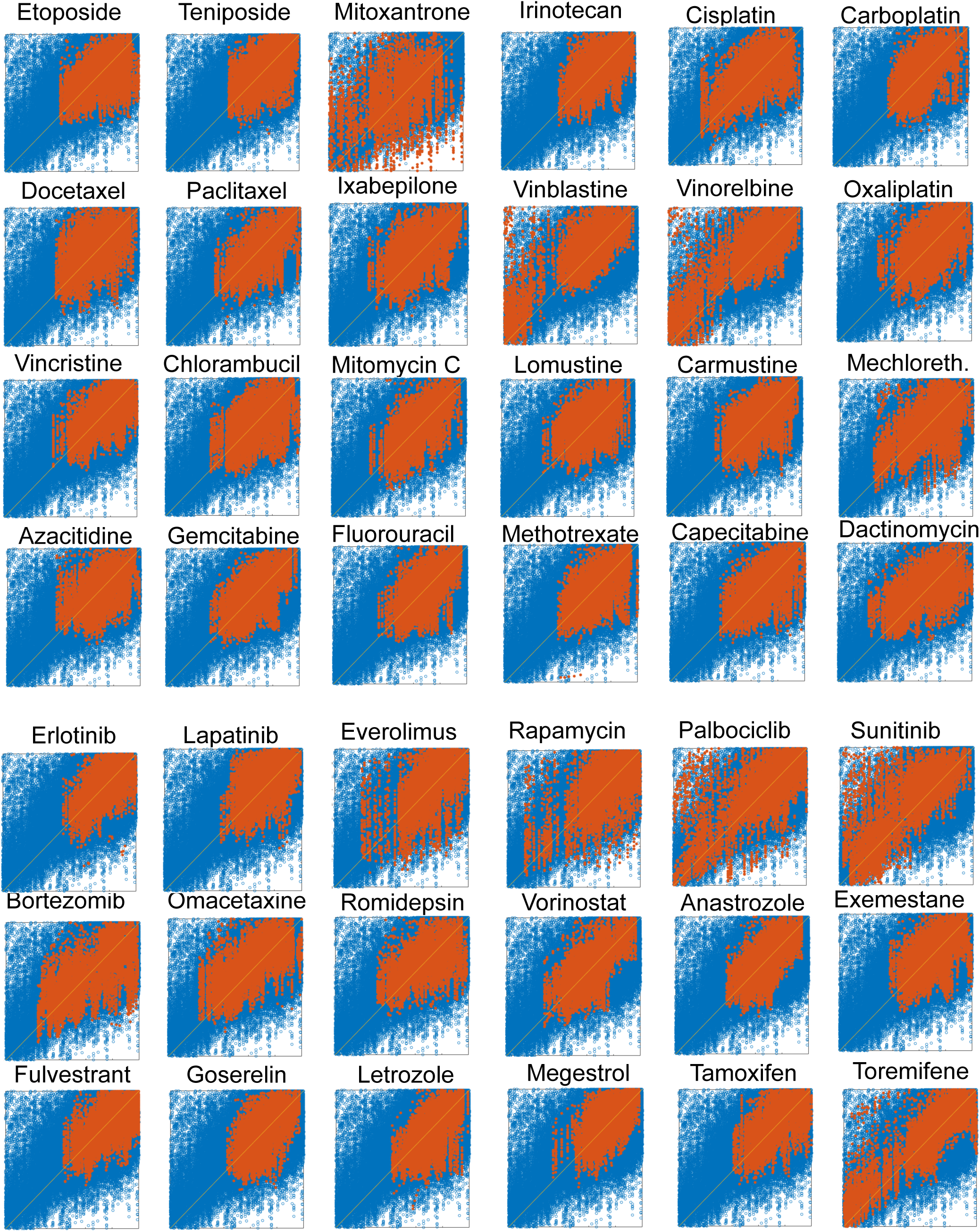
Nearly all classes of drugs are altered by TNBC-CAF interactions. Data plotted as in Figure 2E. Drugs organized by class, with cytotoxic/DNA damaging agents on top (first 4 rows) and targeted therapies below (bottom 3 rows). Data for each drug are highlighted in orange.

**Supplementary Figure 7:**
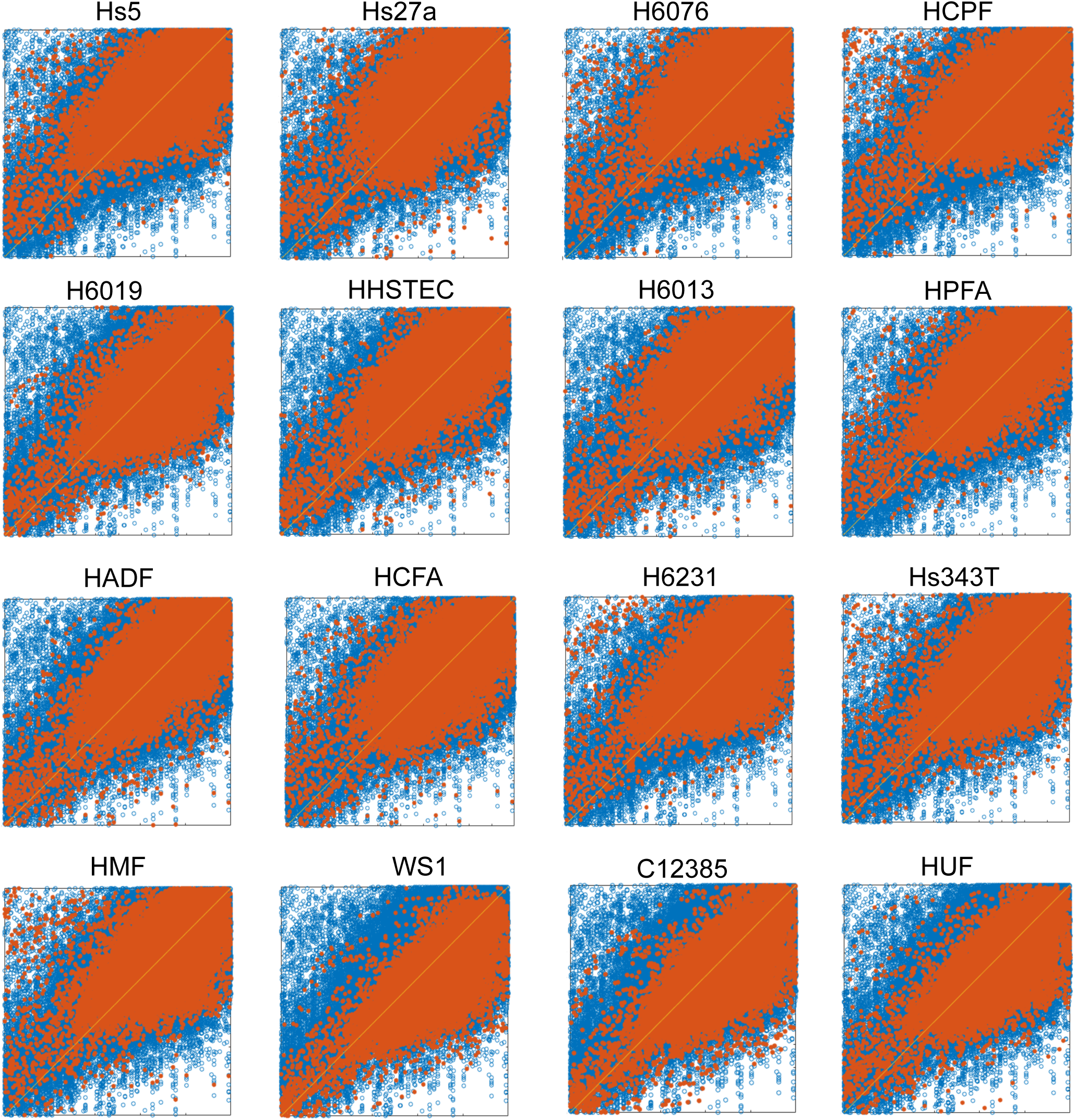
Fibroblasts specific modulation of drug responses in TNBC cells. Data plotted as in Figure 2E. In each plot, the overall dataset is shown in blue circles and the data for each fibroblast cell line are shown in orange. Top 2 rows (8 fibroblast lines) are derived from common metastatic locations; bottom 2 rows are from locations that do not typically harbor breast cancer metastasis.

**Supplementary Figure 8:**
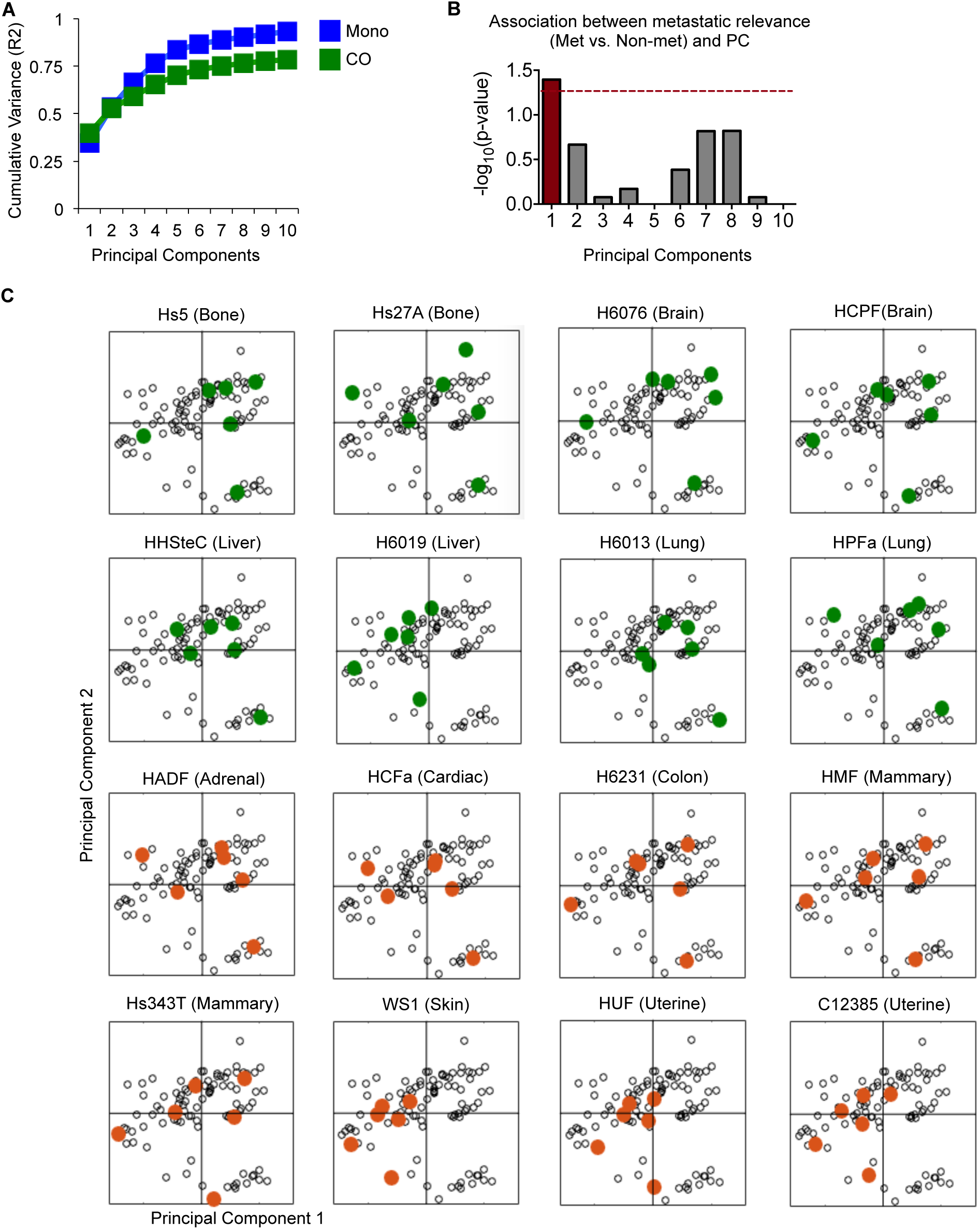
PCA of TNBC drug responses in co-culture reveals divergent roles for CAFs derived from different anatomical locations. **(A)** Cumulative variance (sum of eigenvectors) captured by PCA of mono-culture and co-culture drug responses. **(B)** Statistical association between metastatic class of CAF (common metastatic location or non-common metastatic location) and each principal component. p-values calculated using Fisher’s Exact test. PC1 captures variation associated with met vs. non-met dichotomy. **(C)** PCA scores projection, as in Figure 3A, with scores for each class of CAF highlighted. Common metastatic locations are highlighted in green, and uncommon sites are orange. Positive scores on PC1 are enriched for metastatic sites and negative scores on PC1 are enriched for uncommon sites.

**Supplementary Figure 9:**
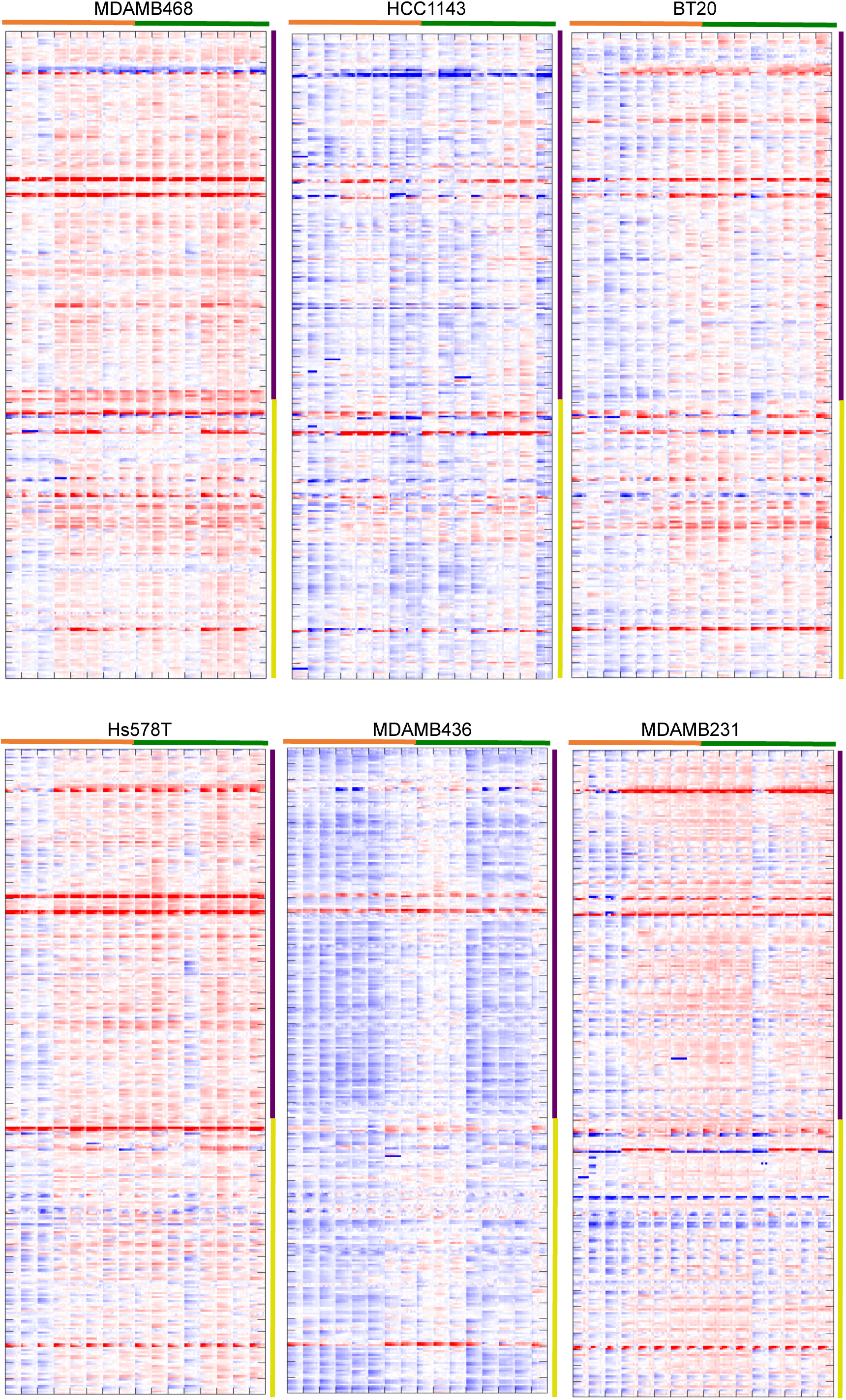
Cell line specific co-culture vs. mono-culture drug response ratios. Data for each cell line are plotted with same order and color scale as in Figure 4B.

## REFERENCES

Al-Lazikani, B., Banerji, U., and Workman, P. (2012). Combinatorial drug therapy for cancer in the post-genomic era. Nat Biotechnol 1–13.

Anders, C., and Carey, L.A. (2008). Understanding and treating triple-negative breast cancer. Oncology (Williston Park, N.Y.) 22, 1233–9–discussion1239–40–1243.

Barretina, J., Caponigro, G., Stransky, N., Venkatesan, K., Margolin, A.A., Kim, S., Wilson, C.J., Lehár, J., Kryukov, G.V., Sonkin, D., et al (2012). The Cancer Cell Line Encyclopedia enables predictive modelling of anticancer drug sensitivity. Nature 483, 603–607.

Cancer Genome Atlas Network (2012). Comprehensive molecular characterization of human colon and rectal cancer. Nature 487, 330–337.

Cancer Genome Atlas Network, Getz, G., Saksena, G., Park, P.J., Chin, L., and Mills, G.B. (2012). Comprehensive molecular portraits of human breast tumours. Nature 490, 61–70.

Carey, L.A., Dees, E.C., Sawyer, L., Gatti, L., Moore, D.T., Collichio, F., Ollila, D.W., Sartor, C.I., Graham, M.L., and Perou, C.M. (2007). The Triple Negative Paradox: Primary Tumor Chemosensitivity of Breast Cancer Subtypes. Clinical Cancer Research 13, 2329–2334.

Chonghaile, T.N., Sarosiek, K.A., Vo, T.T., Ryan, J.A., Tammareddi, A., Moore, V.D.G., Deng, J., Anderson, K.C., Richardson, P., Tai, Y.T., et al (2011). Pretreatment Mitochondrial Priming Correlates with Clinical Response to Cytotoxic Chemotherapy. Science 334, 1129–1133.

Cohen, A.L., Soldi, R., Zhang, H., Gustafson, A.M., Wilcox, R., Welm, B.E., Chang, J.T., Johnson, E., Spira, A., Jeffrey, S.S., et al (2011). A pharmacogenomic method for individualized prediction of drug sensitivity. Mol Syst Biol 7, 1–13.

Fry, R.C., Svensson, J.P., Valiathan, C., Wang, E., Hogan, B.J., Bhattacharya, S., Bugni, J.M., Whittaker, C.A., and Samson, L.D. (2008). Genomic predictors of interindividual differences in response to DNA damaging agents. Genes & Development 22, 2621–2626.

García-González, A.P., Ritter, A.D., Shrestha, S., Andersen, E.C., Yilmaz, L.S., and Walhout, A.J.M. (2017). Bacterial Metabolism Affects the C. elegans Response to Cancer Chemotherapeutics. Cell 169, 431–441.e438.

Gilbert, L.A., and Hemann, M.T. (2010). DNA Damage-Mediated Induction of a Chemoresistant Niche. Cell 143, 355–366.

Heiser, L.M., Wang, N.J., Talcott, C.L., Laderoute, K.R., Knapp, M., Guan, Y., Hu, Z., Ziyad, S., Weber, B.L., Laquerre, S., et al (2009). Integrated analysis of breast cancer cell lines reveals unique signaling pathways. Genome Biol 10, R31.

Innocenti, F., Cox, N.J., and Dolan, M.E. (2011). The use of genomic information to optimize cancer chemotherapy. Semin. Oncol. 38, 186–195.

Janes, K.A., and Yaffe, M.B. (2006). Data-driven modelling of signal-transduction networks. Nat Rev Mol Cell Biol 7, 820–828.

Jiang, T., Shi, W., Wali, V.B., Pongor, L.S., Li, C., Lau, R., Győrffy, B., Lifton, R.P., Symmans, W.F., Pusztai, L., et al (2016). Predictors of Chemosensitivity in Triple Negative Breast Cancer: An Integrated Genomic Analysis. Plos Med 13, e1002193.

Kalluri, R., and Zeisberg, M. (2006). Fibroblasts in cancer. Nat Rev Cancer 6, 392–401.

Lamb, J., Crawford, E.D., Peck, D., Modell, J.W., Blat, I.C., Wrobel, M.J., Lerner, J., Brunet, J.-P., Subramanian, A., Ross, K.N., et al (2006). The Connectivity Map: using gene-expression signatures to connect small molecules, genes, and disease. Science 313, 1929–1935.

Lamprecht, M., Sabatini, D., and Carpenter, A. (2007). CellProfiler(tm): free, versatile software for automated biological image analysis. Biotech. 42, 71–75.

Lee, M.J., Ye, A.S., Gardino, A.K., Heijink, A.M., Sorger, P.K., Macbeath, G., and Yaffe, M.B. (2012). Sequential application of anticancer drugs enhances cell death by rewiring apoptotic signaling networks. Cell 149, 780–794.

Lehmann, B.D., Bauer, J.A., Chen, X., Sanders, M.E., Chakravarthy, A.B., Shyr, Y., and Pietenpol, J.A. (2011). Identification of human triple-negative breast cancer subtypes and preclinical models for selection of targeted therapies. J Clin Invest 1–18.

Li, J., Zhao, W., Akbani, R., Liu, W., Ju, Z., Ling, S., Vellano, C.P., Roebuck, P., Yu, Q., Eterovic, A.K., et al (2017). Characterization of Human Cancer Cell Lines by Reverse-phase Protein Arrays. Cancer Cell 31, 225–239.

Mcmillin, D.W., Delmore, J., Weisberg, E., Negri, J.M., Geer, D.C., Klippel, S., Mitsiades, N., Schlossman, R.L., Munshi, N.C., Kung, A.L., et al (2010). Tumor cell-specific bioluminescence platform to identify stroma-induced changes to anticancer drug activity. Nat Med 16, 483–489.

Montero, J., Sarosiek, K.A., DeAngelo, J.D., Maertens, O., Ryan, J., Ercan, D., Piao, H., Horowitz, N.S., Berkowitz, R.S., Matulonis, U., et al (2015). Drug-induced death signaling strategy rapidly predicts cancer response to chemotherapy. Cell 160, 977–989.

Nguyen, T.V., Sleiman, M., Moriarty, T., Herrick, W.G., and Peyton, S.R. (2014). Sorafenib resistance and JNK signaling in carcinoma during extracellular matrix stiffening. Biomaterials 35, 5749–5759.

Pallasch, C.P., Leskov, I., Braun, C.J., Vorholt, D., Drake, A., Soto-Feliciano, Y.M., Bent, E.H., Schwamb, J., Iliopoulou, B., Kutsch, N., et al (2014). Sensitizing Protective Tumor Microenvironments to Antibody-Mediated Therapy. Cell 156, 590–602.

Perou, C.M., Sørlie, T., Eisen, M.B., van de Rijn, M., Jeffrey, S.S., Rees, C.A., Pollack, J.R., Ross, D.T., Johnsen, H., Akslen, L.A., et al (2000). Molecular portraits of human breast tumours. Nature 406, 747–752.

Prat, A., Parker, J.S., Karginova, O., Fan, C., Livasy, C., Herschkowitz, J.I., He, X., and Perou, C.M. (2010). Phenotypic and molecular characterization of the claudin-low intrinsic subtype of breast cancer. Breast Cancer Res 12, R68–18.

Ryan, J., and Letai, A. (2013). BH3 profiling in whole cells by fluorimeter or FACS. Methods 61, 156–164.

Ryan, J., Montero, J., Rocco, J., and Letai, A. iBH3: simple, fixable BH3 profiling to determine apoptotic priming in primary tissue by flow cytometry. Biological Chemistry 397, 14.

Shah, S.P., Roth, A., Goya, R., Oloumi, A., Ha, G., Zhao, Y., Turashvili, G., Ding, J., Tse, K., Haffari, G., et al (2012). The clonal and mutational evolution spectrum of primary triple-negative breast cancers. Nature 1–5.

Straussman, R., Morikawa, T., Shee, K., Barzily-Rokni, M., Qian, Z.R., Du, J., Davis, A., Mongare, M.M., Gould, J., Frederick, D.T., et al (2012). Tumour micro-environment elicits innate resistance to RAF inhibitors through HGF secretion. Nature 487, 500–504.

Weaver, V.M., Lelièvre, S., Lakins, J.N., Chrenek, M.A., Jones, J.C.R., Giancotti, F., Werb, Z., and Bissell, M.J. (2002). beta4 integrin-dependent formation of polarized three-dimensional architecture confers resistance to apoptosis in normal and malignant mammary epithelium. Cancer Cell 2, 205–216.

Yard, B.D., Adams, D.J., Chie, E.K., Tamayo, P., Battaglia, J.S., Gopal, P., Rogacki, K., Pearson, B.E., Phillips, J., Raymond, D.P., et al (2016). A genetic basis for the variation in the vulnerability of cancer to DNA damage. Nature Communications 7, 11428.

